# GATAD2B containing NuRD complex drives R-loop dependent chromatin boundary formation at double strand breaks

**DOI:** 10.1101/2024.02.19.581003

**Authors:** Zhichao Liu, Kamal Ajit, Yupei Wu, Wei-Guo Zhu, Monika Gullerova

## Abstract

Double-strand breaks (DSBs) are the most lethal form of DNA damage. Transcriptional activity at DSBs, as well as transcriptional repression around DSBs, are both required for efficient DNA repair. The chromatin landscape defines and coordinates these two opposing events. However, the regulation of the open and condensed chromatin architecture is still unclear. In this study, we show that the GATAD2B-NuRD complex associates with DSBs in a transcription- and R-loop-dependent manner, to promote histone deacetylation and chromatin condensation, creating a temporal boundary between open and closed chromatin. This boundary is necessary for correct DNA end resection termination. The lack of the GATAD2B-NuRD complex leads to chromatin hyper-relaxation and extended DNA end resection, resulting in HR repair failure. Our results suggest that the GATAD2B-NuRD complex is a key coordinator of the dynamic interplay between transcription and chromatin landscape and underscore its biological significance in the RNA-dependent DNA damage response.

## Introduction

Maintenance of genome stability is crucial for normal biological and cellular activities(Jackson & Bartek, 2009). The human genome is continuously exposed to various external or internal factors that could lead to different types of DNA damage, of which DNA double strand breaks (DSBs) are the most lethal. Fatal consequences, including loss of genetic information, cell death, mutagenesis and premature aging, could be triggered, if DNA damage is left unrepaired or repaired incorrectly(McKinnon, 2009; White & Vijg, 2016). To recognize, signal and correctly repair DNA breaks, cells have developed a complex pathway called the DNA damage response (DDR)(Ciccia & Elledge, 2010). The majority of DSBs will be repaired by two major pathways, homologous recombination (HR) or non-homologous end joining (NHEJ). The canonical and precise HR pathway utilizes sister chromatids as the DNA template. HR employs DNA end resection to create a single-stranded DNA (ssDNA) overhang, which is in turn used to search for and anneal with the intact DNA template of the sister chromatid(Heyer *et al*, 2010). On the other hand, error-prone NHEJ may rapidly ligate two broken DNA ends(Lieber, 2010) in any stage of the cell cycle.

Recently, several studies have revealed that transcription at DSBs plays an important role in DDR (Aymard *et al*, 2014; Ketley *et al*, 2022; Ketley & Gullerova, 2020; Long *et al*, 2021). Specifically, phosphorylated RNA polymerase II (RNAPII) transcribes long non-coding RNA, such as damage-induced lncRNA (dilncRNA)(Michelini *et al*, 2017; Pessina *et al*, 2019) and damage-responsive transcripts (DARTs)(Burger *et al*, 2019). DSB-derived long non-coding RNAs have been shown to be further processed into small non-coding RNAs named DNA-damage derived RNAs (DDRNAs)(Bonath *et al*, 2018; Francia *et al*, 2012) and were suggested to facilitate the recruitment of DDR factors, including 53BP1, to DSBs. DilncRNA can also anneal with ssDNA overhang template after resection to form DNA:RNA hybrids(D’Alessandro *et al*, 2018; Francia *et al*, 2016). DNA:RNA hybrids, are widely considered to be a transcription byproduct and a potential threat to genomic stability(Skourti-Stathaki & Proudfoot, 2014). However, they are also found to be formed at DSB sites predominantly in transcriptionally active loci(Bader & Bushell, 2020; Cohen *et al*, 2018; Marnef & Legube, 2021; Yasuhara *et al*, 2018). R-loops (DNA:RNA hybrids together with the ssDNA strand) formed behind pausing RNAPII, were implicated in regulation of repair pathway choices and recruitment of DSB factors, such as CSB, RPA, Rad52, BRCA1, BRCA2 and 53BP1(Burger *et al*., 2019; D’Alessandro *et al*., 2018; Hatchi *et al*, 2015; Mazina *et al*, 2020; Tan *et al*, 2020; Teng *et al*, 2018; Yasuhara *et al*., 2018). R-loops are involved in regulation of several biological processes under normal conditions(Garcia-Muse & Aguilera, 2019; Petermann *et al*, 2022). Specifically, they are associated with active transcription at promoter regions(Ginno *et al*, 2012; Santos-Pereira & Aguilera, 2015), facilitate transcription termination(Skourti-Stathaki *et al*, 2014) and regulate chromatin modification and remodeling(Castellano-Pozo *et al*, 2013; Herrera-Moyano *et al*, 2014; Prendergast *et al*, 2020). Whether R-loops or DNA:RNA hybrids play these or similar roles at DSBs remains enigmatic.

Chromatin architecture is regulated by chromatin remodeling complexes and histone modifying enzymes. Generally open chromatin is associated with active transcription, whilst condensed heterochromatin represents transcriptionally silent regions. Remodeling of chromatin landscape is an important step in DDR(Price & D’Andrea, 2013). Several studies showed that upon DNA damage, chromatin opens up to promote transcription, the loading of DDR factors and efficient completion of DNA repair (Hauer *et al*, 2017; Luijsterburg *et al*, 2016; Murr *et al*, 2006). On the other hand, several proteins involved in chromatin compaction and heterochromatin formation were detected around DSBs(Ayrapetov *et al*, 2014; Kalousi *et al*, 2015; Yang *et al*, 2017). Furthermore, recent bioimaging data detected opened chromatin at DSBs, which was surrounded by compacted chromatin(Lou *et al*, 2019), suggesting that there might be a border between these two types of chromatin landscapes. However, the mechanism and dynamics of chromatin structure and function upon DNA damage remains unclear.

Here, we employed mass spectroscopy-based proteomics to screen proteins, which bind to R-loops upon DNA damage and identified GATAD2B and MBD3, two subunits of Nucleosome Remodeling Deacetylase (NuRD) complex, among significant hits. Both, GATAD2B and MBD3 are associated with DSBs in a transcriptional and R-loop dependent manner. At damaged loci, GATAD2B-NuRD complex promotes histone deacetylation followed by chromatin condensation. Lack of GATAD2B leads to chromatin hyper-relaxation and the consequently to DNA end hyper-resection and failed HR repair. Overall, our results reveal that transcription and R-loops recruit GATAD2B-NuRD complex to establish temporal boundaries between open and condensed chromatin around DSBs. Chromatin boundaries act to promote termination of the end resection and to facilitate efficient HR repair.

## Results

### Components of NuRD complex bind to R-loops upon DNA damage

To identify novel R-loop interactors relevant to DDR, we employed the R-loop specific S9.6 antibody (Cristini *et al*, 2018) for mass spectrometry-based proteomics (Fig 1A). The HEK293T cells were subjected to 10 Gy irradiation (IR) treatment and followed by 20 min recovery. First, we pulled down proteins bound to S9.6 antibody and tested, whether we can detect known R-loop interactors in our IP fractions. Indeed, we detected previously identified R-loop interactors such as DHX9 and XRN2 in our IP samples (Figs EV1A-E)(Cristini *et al*., 2018). Next, we performed S9.6 immunoprecipitation from control (-IR) and irradiated (+IR) HEK293 cells, followed by mass spectrometry-based proteomics. In our control (-IR) samples we identified many well-known R-loops binders, such as DDX5, XRN2 or PARP1(Laspata *et al*, 2023) (Fig EV1F). We also identified proteins, which were significantly enriched in S9.6 IP samples specifically upon DNA damage (Figs 1B and Fig EV1G). GO and STRING analyses revealed enrichment for proteins involved in RNA processing, gene expression regulation and histone deacetylation (Fig 1C and D). Interestingly, we found MBD3 and GATAD2B, two components of the NuRD complex, among damage specific R-loop interactors(Torchy *et al*, 2015). NuRD complex has histone deacetylation as well as ATP-dependent chromatin remodeling subunits, and has been found to regulate several biological processes, including transcription repression(Torchy *et al*., 2015).

**Fig 1:**
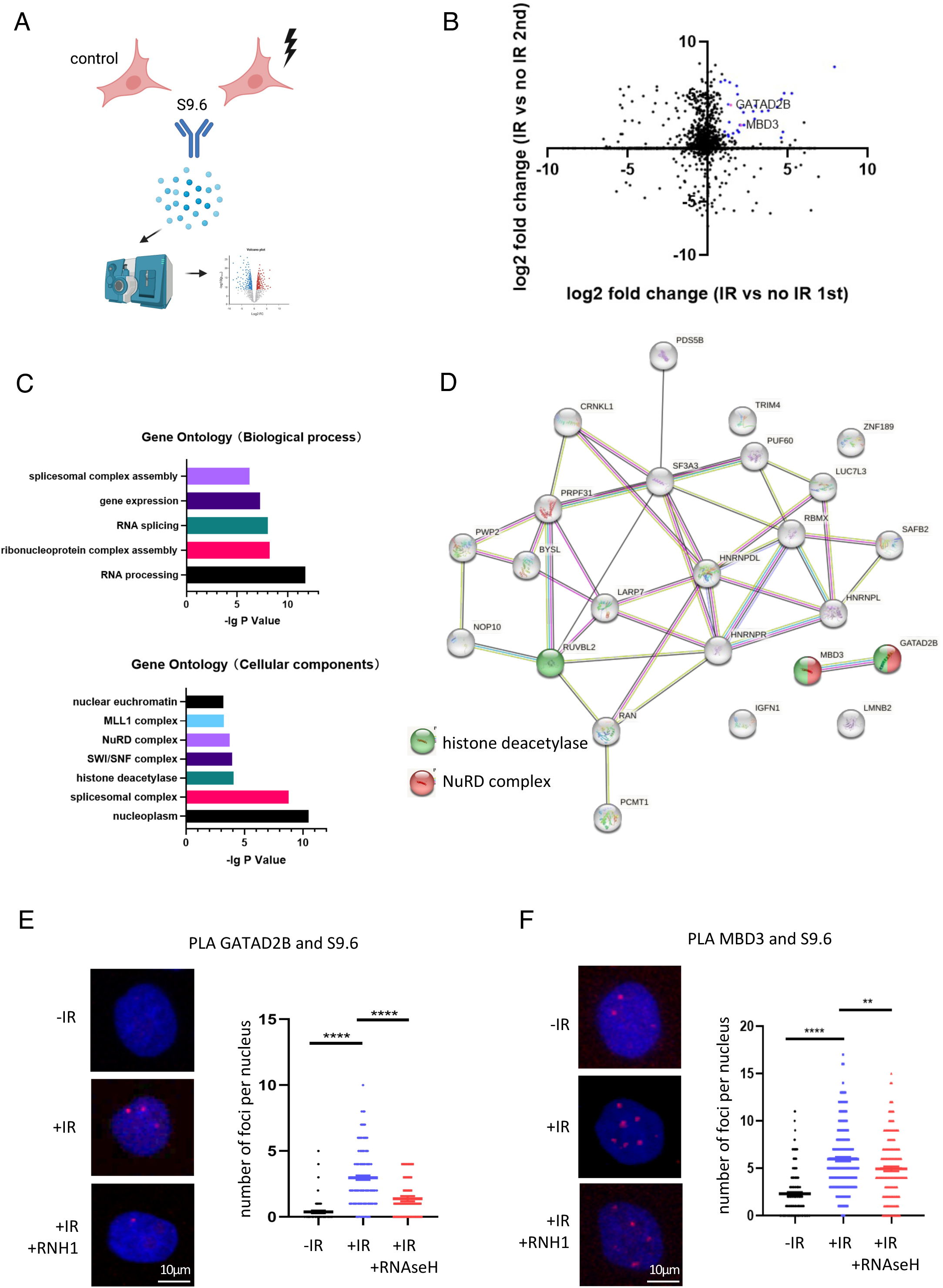
Components of NuRD complex bind to R-loops upon DNA damage. A) Diagram of S9.6 mass spectroscopy-based proteomics approach. B) Volcano plot showing significantly enriched S9.6 binders after IR treatment in two biological replicates. GATAD2B and MBD3 are highlighted and named. C) GO analysis of proteins significantly enriched in S9.6 IP after IR treatment. D) STRING analysis of proteins significantly enriched in S9.6 IP after IR treatment. E) PLA of GATAD2B and S9.6 in cells with or without IR and overexpression of RNAseH1. IR=10Gy. Left: representative confocal microscopy images; right: quantification of left, error bar = mean ± SEM, significance was determined using non-parametric Mann-Whitney test. ****p ≤ 0.0001. F) PLA of MBD3 and S9.6 in cells with or without IR and overexpression of RNAseH1. IR=10Gy. Left: representative confocal microscopy images; right: quantification of left, error bar = mean ± SEM, significance was determined using non-parametric Mann-Whitney test. ***p ≤ 0.001, \*\**p* ≤ 0.01.

To further validate whether components of NuRD complex could interact with R-loops upon DNA damage *in vivo*, we employed optimized proximity ligation assay (PLA) that allows for detection of protein/R-loop complexes *in vivo*(Alagia *et al*, 2022). Using antibodies against endogenous GATAD2B or MBD3 and S9.6 we detected significant increase in PLA foci upon IR treatment. Number of PLA foci was decreased in presence of RNAseH1 (enzyme resolving R-loops), which confirms the specificity of this experimental approach (Figs 1E and 1F). Single antibodies were used as PLA negative controls (Fig EV2A). To test whether binding of GATAD2B to R-loops is dependent on any of the DDR signaling pathways such as ATM or PARP1, we performed GATAD2B and S9.6 PLA in cells treated with ATM inhibitor or PARP1 inhibitors and detected that inhibition of PARP1 significantly decreased binding of GATAD2B to R-loops, whilst inhibition of ATM did not (Fig EV2B). Histone deacetylation activity of NuRD complex is facilitated by histone deacetylase 1, HDAC1, subunit. To test whether HDAC1 could also bind to R-loops, we performed PLA using S9.6 and HDAC1 antibodies and detected significantly increased number of RNAseH1 sensitive PLA foci upon IR treatment (Fig EV2C).

These results show that the subunits of GATAD2B-NuRD complex preferentially bind to R-loops upon DNA damage in PARP1 dependent manner.

### GATAD2B-NuRD complex are recruited to DSBs in transcription and R-loop dependent manner

The above data identified components of NuRD complex as a novel DNA damage specific R-loop interactor. R-loops are associated with transcription and transcription at DSBs is required for efficient DNA repair(Yasuhara *et al*., 2018). Therefore, we wished to test whether the GATAD2B-NuRD complex is recruited to DSBs in an R-loop dependent manner. HeLa cells were transfected with GFP-GATAD2B and GFP-MBD3 cells respectively, and subjected to laser micro-irradiation followed by time-lapse microscopy. We found that both proteins were rapidly recruited to laser-induced DNA damage stripes in less than 30s. Interestingly, upon transcription inhibition by triptolide or DRB, recruitment of both GATAD2B and MBD3 to DSBs was significantly reduced (Figs 2A and Fig EV3A). Next, we transfected cells with RNaseH1-mCherry and performed laser stripping. Live cell imaging revealed that localization of GATAD2B and MBD3 at DSBs was reduced by the overexpression of RNaseH1 (Fig 2B and Fig EV3B). Furthermore, kinetics of both, RNaseH1 and GATAD2B recruitment to DSBs, showed similar profile, suggesting that there might be a correlation in their binding to DSBs (Fig EV3C).

**Fig 2.**
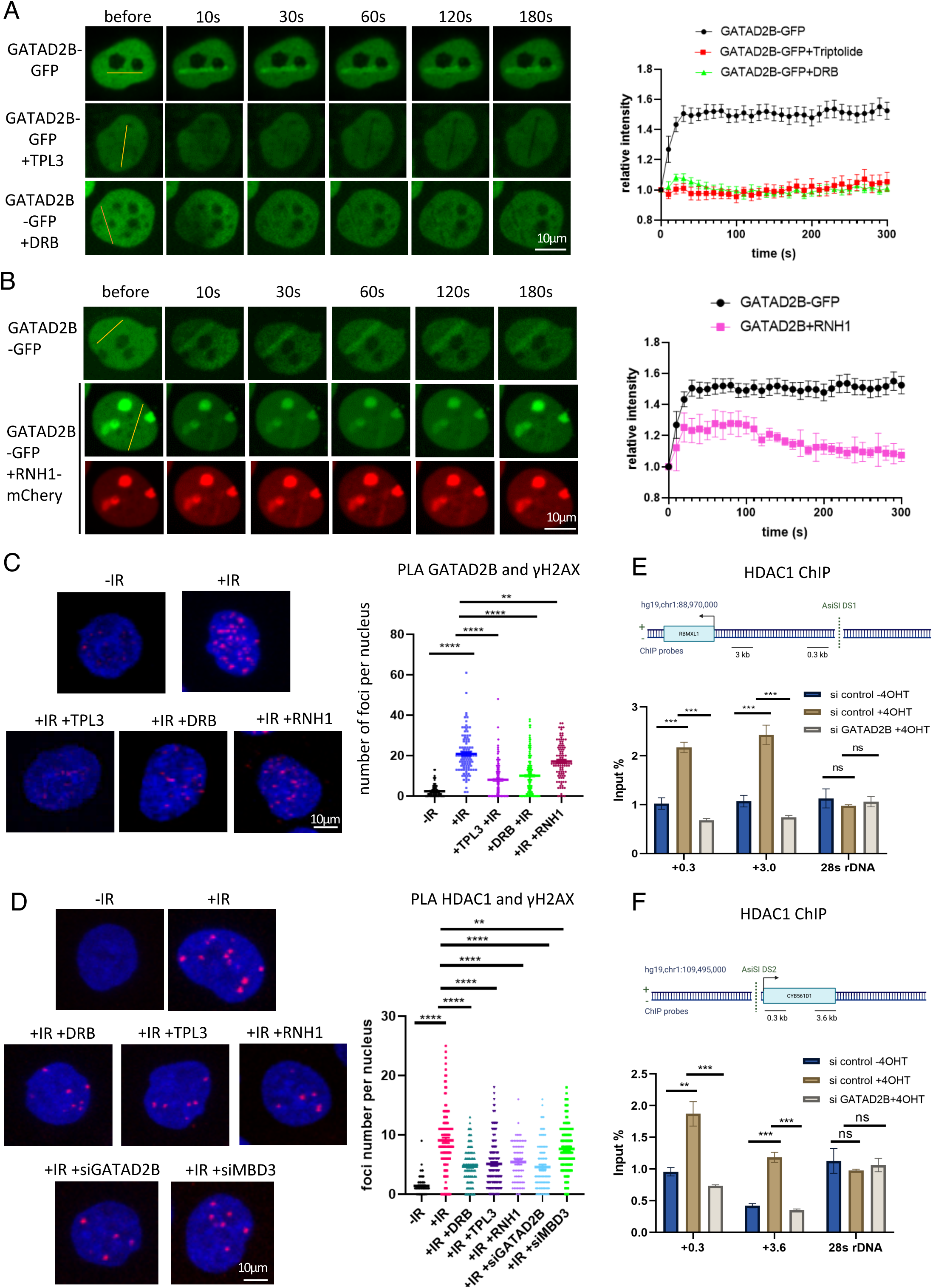
GATAD2B is recruited to DSBs in transcription and R-loop dependent manner and required for HDAC1 localization to DSBs. A) Laser stripping of GATAD2B-GFP cells with or without treatment with transcription inhibitors triptolite (TPL3) and DRB. Representative spinning disk confocal microscopy images and quantification (n≥10) showing GFP signals before and after laser striping at indicated time points; error bar = mean ± SEM. B) Laser stripping of GATAD2B-GFP cells with or without transiently expression of RNAseH1-mCherry plasmid. Representative spinning disk confocal microscopy images and quantification (n≥10) showing GFP and RFP signals before and after laser striping at indicated time points; error bar = mean ± SEM. C) PLA of GATAD2B and γH2AX in cells with or without IR and transcription inhibition (TLP3 or DRB) or overexpression of RNAseH1. IR=10Gy. Left: representative confocal microscopy images; right: quantification of left, error bar = mean ± SEM, significance was determined using non-parametric Mann-Whitney test. ****p ≤ 0.0001, \*\**p* ≤ 0.01. D) PLA of HDAC1 and γH2AX in cells with or without IR and transcription inhibition (TLP3 or DRB), overexpression of RNAseH1 or depletion of GATAD2B and MBD3. IR=10Gy. Left: representative confocal microscopy images; right: quantification of left, error bar = mean ± SEM, significance was determined using non-parametric Mann-Whitney test. ****p ≤ 0.0001, \*\**p* ≤ 0.01. E) Top: drawing showing DS1 genomic region with AsiSI cut site and position of ChIP probes. Bottom: Bar chart showing HDAC1 ChIP levels at indicated sites next to AsiSI cut in cells with or without 4OH and depletion of GATAD2B, error bar = mean ± SEM, significance was determined using non-parametric Mann-Whitney test. ***p ≤ 0.001, n.s. not significant. F) As in E, for DS2.

Next, we investigated whether the interaction between R-loops and GATAD2B is specifically regulated as a part of DDR or whether DNA damage causes retention of NuRD at actively transcribed regions, to those it is known to bind. We selected three genes known to be bound by NuRD complex and performed chromatin immunoprecipitation in cells overexpressing RNaseH1 or treated with transcription inhibitor (Triptolite). We show that neither transcription inhibition nor RNAseH1 overexpression affect the level of NuRD at these loci in non-damage condition. These data suggest that NuRD complex is recruited de novo to DSBs in R-loop and transcription dependent manner (Fig EV3D).

To further support these results, we performed PLA using GATAD2B or MBD3 and γH2AX antibodies and show increased number of PLA foci upon IR treatment when compared to control cells. Both, GATAD2B/γH2AX and MBD3/γH2AX PLA foci were sensitive to transcription inhibition (TPL3 and DRB treatment) and R-loop removal (overexpression of RNAseH1) (Fig 2C and Fig EV4A, single antibodies were used as PLA negative controls).

Next, we wished to test whether GATAD2B as a component of NuRD complex, plays a role in HDAC1 localisation to DSBs. We performed PLA using HDAC1 and γH2AX antibodies and detected PLA foci upon IR treatment. Furthermore, these PLA foci were sensitive to transcription inhibition (DRB and TPL3), as well as RNAseH1 overexpression. Interestingly, depletion of GATAD2B also resulted in reduced number of HDAC1/ γH2AX PLA foci (Fig 2D and Fig EV4B). To support these data, we also performed HDAC1 ChIP in the DIvA cell line, where ER-fused AsiSI restriction enzyme can be translocated into nucleus upon addition of tamoxifen (4OH) and cleave DNA at specific sites, simulating double strand breaks. This cellular system is commonly used to study DSBs in sequence specific manner (Aymard *et al*., 2014; Iacovoni *et al*, 2010). Indeed, we detected HDAC1 ChIP signal across two different AsiSI cut sites, when damage was induced and the levels of HDAC1 were significantly decreased by GATAD2B depletion (Fig 2E and F).

KDM5A and ZMYND8 are two histone modifiers that play important roles in the DDR by interacting with PAR and recruiting the NuRD complex to DNA damage sites. KDM5A is a histone demethylase that binds to PAR chains through a noncanonical region and represses transcription to facilitate HR repair(Kumbhar *et al*, 2021). ZMYND8 is a BRD protein that recognizes acetylated chromatin and mediates a transcription-associated DDR pathway that targets actively transcribed regions. ZMYND8 also controls PAR levels at DNA breaks by recruiting ARH3 and modulates the recruitment of BRCA1 and RAD51, two key factors for HR repair (Gong *et al*, 2015; Gong *et al*, 2017; Kumbhar *et al*., 2021; Spruijt *et al*, 2016). Therefore, next we tested whether these two proteins might be involved in recruitment of GATAD2B-NuRD to DSBs. PLA using antibodies against HDAC1 and γH2AX in cells depleted of MBD3, GATAD2B, ZMYND8 and KDM5A, revealed that ZMYND8 KD, as well as GATAD2B and MBD3 KD, significantly reduced number of PLA foci (Fig EV4C).

Our results suggest that GATAD2B-NuRD complex is recruited to DSBs in transcription and R-loop dependent manner and is facilitating HDAC1 recruitment to DSBs through GATAD2B. ZMYND8 is also required for HDAC1 recruitment to DSBs most likely through its role in transcription associated DDR and binding to PAR.

### GATAD2B and HDAC1 associate with transcriptionally active and HR prone DSBs

To elucidate the genome wide distribution and profile of GATAD2B-NuRD complex around DSBs, we employed DIvA cell line(Aymard *et al*., 2014; Iacovoni *et al*., 2010) and performed chromatin immunoprecipitation using antibodies against GATAD2B and HDAC1, followed by next generation sequencing (ChIP-seq). All three replicates of ChIP-seq data were assessed by Principal Component Analysis (PCA) and show the reproducibility of the data (Fig EV5A).

DSBs (as cut sites) were identified and mapped by BLESS technique(Clouaire *et al*, 2018a). ChIP-seq analysis showed enrichment of both GATAD2B and HDAC1 at DSBs, when cut was induced by 4OHT treatment (Figs 3A-D and Figs EV5B, EV6A and B; levels of input signals were used as a control). The accumulation of both proteins was most prominent at DSBs (as defined by BLESS signal) and spreads across in a 2.5-kb window around AsiSI cutting sites. Next, we overlapped our GATAD2B and HDAC1 ChIP-sep profiles with DRIP-seq data showing the presence of R-loops near DSBs(Cohen *et al*., 2018). DRIP-seq technique employs S9.6 antibody pull down followed by DNA sequencing. As the DNA at the breaks is being cut, there are no detectable DRIP signals right at the breaks. Hence DRIP-seq metagene profiles results in dip in sequencing reads at DSBs. This is not the case for ChIP-seq approach. Overlay of our ChIP-seq and DRIP-seq metagene profiles revealed that R-loops were surrounding and partially overlapping the GATAD2B, HDAC1 peaks. Interestingly, all metagene profiles showed drop in intensity at around 2.5 kb away from the breaks (Fig 3A). Heatmap analysis revealed that the recruitment of GATAD2B-NuRD complex correlated with AsiSI cutting efficiency, indicating that this complex was prone to be recruited to more accessible AsiSI sites, likely to be near transcriptionally active regions (Fig EV6B). Indeed, both, GATAD2B and HDAC1, were more enriched at transcription active loci, when compared to transcriptionally repressed regions (Figs 3E and F, Fig EV7A-C). Previous reports have shown that transcription and R-loops could serve as a binding platform for recruitment of HR repair pathway proteins(Aymard *et al*., 2014). AsiSI cut sites in DIvA system can be divided into HR or NHEJ prone DSBs based on correlation ratio using ChIP-Seq data coverage of RAD51, well known facilitator of HR, (AsiSI site +- 4kb) to XRCC4, known NHEJ factor, coverage (AsiSI site +- 1kb). Top 30 sites were annotated as HR prone due to higher coverage of RAD51, and the bottom 30 as NHEJ prone due to higher ratio of XRCC4(Clouaire *et al*., 2018a). ChIP-seq data revealed that both GATAD2B and HDAC1 are significantly more associated with HR prone AsiSI cut sites (Figs 3G and H and Fig EV7D-F).

**Fig 3.**
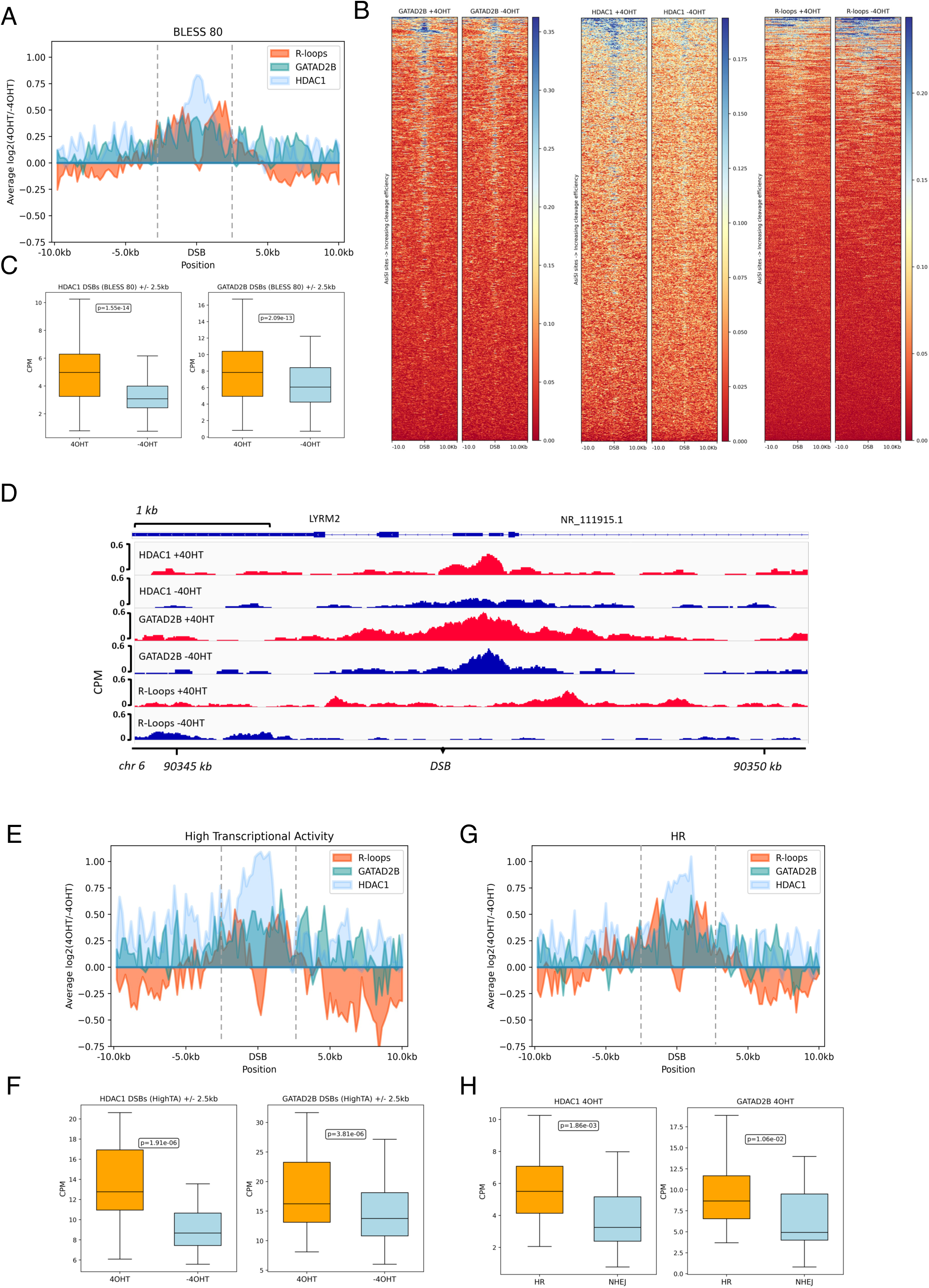
GATAD2B and HDAC1 form boundary around DSBs in transcriptionally active and HR prone loci. A) Metagene profile showing ChIP-seq enrichment of GATAD2B, HDAC1 and R-loops at AsiSI cut sites (as defined by BLESS technique). B) Heatmaps showing GATAD2B and HDAC1 ChIP-seq and DRIP-seq read count over a 20 kb window centered on the DSB before (−4OHT) and after (+4OHT) DSB induction. DSBs are sorted according to decreasing cleavage efficiency, as defined by BLESS technique. See text for references. C) Box plot graph showing levels of HDAC1 and GATAD2B ChIP-seq reads mapping to AsiSI sites in presence or absence of 4OHT. D) IGV screenshot showing GATAD2B and MBD3 ChIP-Seq and DRIP-seq reads before (−4OHT) and after damage induction (+4OHT) at selected AsiSI site. E) Metagene profile showing ChIP-seq enrichment of GATAD2B, HDAC1 and R-loops at AsiSI cut sites in highly transcribed regions. F) Box plot graph showing levels of HDAC1 and GATAD2B ChIP-seq reads mapping to AsiSI sites in loci with high or low transcriptional activity. G) Metagene profile showing ChIP-seq enrichment of GATAD2B, HDAC1 and R-loops at HR prone AsiSI cut sites. H) Box plot graph showing levels of HDAC1 and GATAD2B ChIP-seq reads mapping to AsiSI sites in HR- or NHEJ-prone loci.

Overall, our genome wide data show that the GATAD2B-NuRD complex is mostly prominent at and surrounded by R-loops around DSBs, suggesting that R-loops form a platform for NuRD recruitment to DSBs. The GATAD2B-NuRD complex is more enriched at HR-prone DSBs and regions with a higher transcription activity.

### GATAD2B-NuRD complex promotes histone deacetylation around DSBs

We show above that GATAD2B binding to R-loops is required for NuRD complex localization at DSBs. As NuRD complex contains histone deacetylase HDAC1, we examined next, whether it negatively correlates with histone acetylation levels around DSBs. Acetylation of histone H4 lysine residue is associated with transcriptionally active genomic loci. Interestingly, the ChIP-seq profile of acetylated H4K12 (H4K12Ac) in DIvA cells shows the presence of this histone modification at uncut AsiSI sites, but at reduced levels following cut induction (Fig EV8A). The alignment of our GATAD2B and HDAC1 ChIP-seq data with the H4K12Ac ChIP-seq profile (Clouaire *et al*., 2018a) revealed a negative correlation between H4K12Ac levels and GATAD2B or HDAC1 levels around DSBs (Fig 4A-C and Fig EV8A and EV9A-C). These opposite profiles were more pronounced at the sites with high transcription activity and HR prone DSBs (Fig 4D, E and Fig EV8B and C).

**Fig 4.**
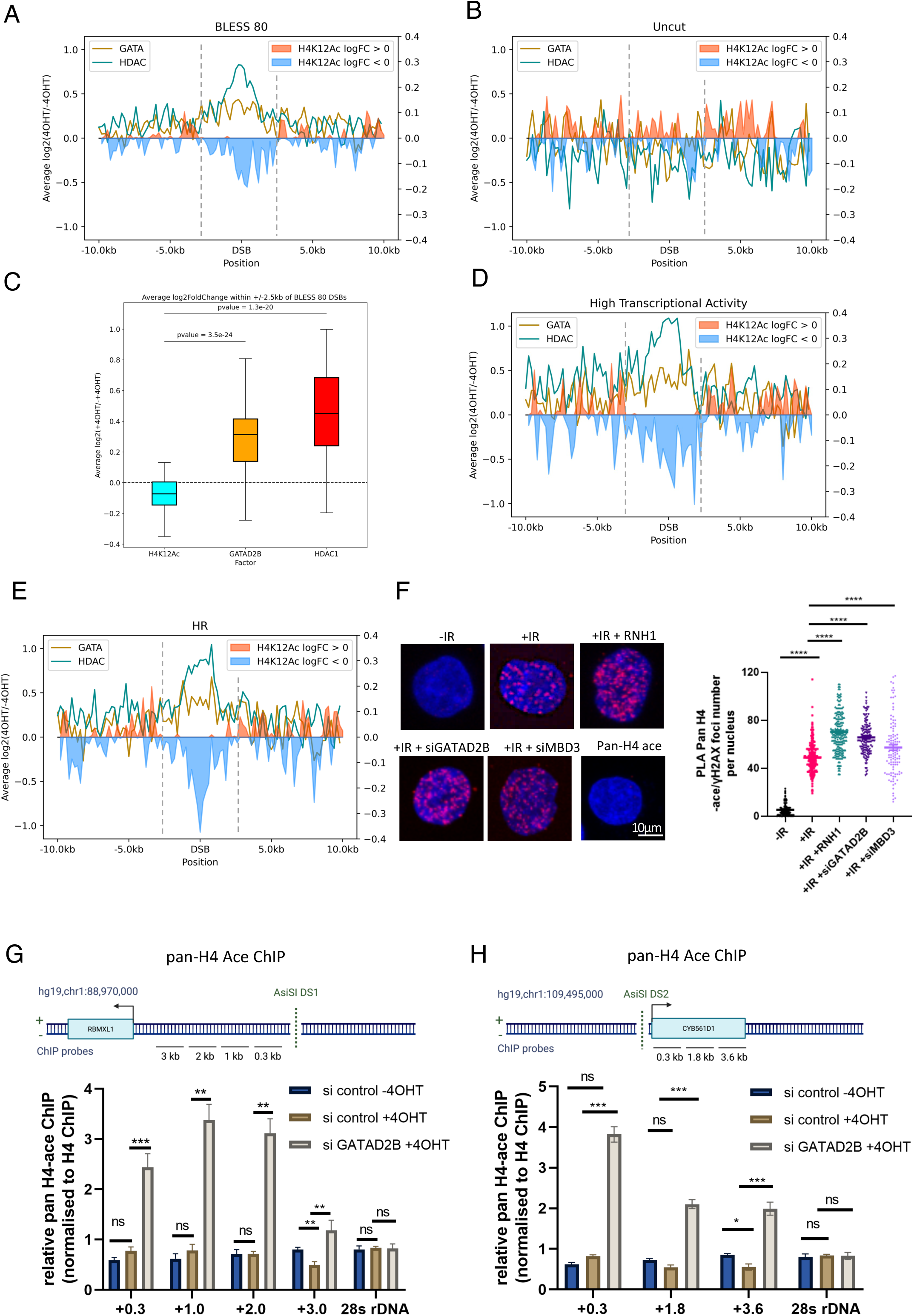
GATAD2B-NuRD complex promotes histone de-acetylation at DSBs. A) Metagene profile showing ChIP-seq enrichment of GATAD2B, HDAC1 and H4K12ac at cut AsiSI (as defined by BLESS technique). B) Metagene profile showing ChIP-seq enrichment of GATAD2B, HDAC1 and H4K12ac at uncut AsiSI sites (as defined by BLESS technique). C) Box plot showing average log2FoldChange within 2,5 kb around cut AsiSI for H4K12ac, GATAD2B and HDAC1 ChIP-seq counts. D) Metagene profile showing ChIP-seq enrichment of GATAD2B, HDAC1 and H4K12ac at cut AsiSI sites in highly transcribed regions. E) Metagene profile showing ChIP-seq enrichment of GATAD2B, HDAC1 and H4K12ac at HR prone cut AsiSI sites. F) PLA of pan-acetyl H4 and γH2AX in cells with or without IR, and overexpression of RNAseH1 or depletion of GATAD2B and MBD3. IR=10Gy. Left: representative confocal microscopy images; right: quantification of left, error bar = mean ± SEM, significance was determined using non-parametric Mann-Whitney test. ****p ≤ 0.0001 G) Top: drawing showing DS1 genomic region with AsiSI cut site and position of ChIP-qPCR probes. Bottom: Bar chart showing relative pan-acetyl H4 ChIP levels (normalised to H4 ChIP levels) at indicated sites next to AsiSI cut in cells with or without 4OH and depletion of GATAD2B, error bar = mean ± SEM, significance was determined using non-parametric Mann-Whitney test. ***p ≤ 0.001, **p ≤ 0.01, n.s. not significant. H) Top: drawing showing DS2 genomic region with AsiSI cut site and position of ChIP-qPCR probes. Bottom: Bar chart showing relative pan-acetyl H4 ChIP levels (normalised to H4 ChIP) at indicated sites next to AsiSI cut in cells with or without 4OH and depletion of GATAD2B, error bar = mean ± SEM, significance was determined using non-parametric Mann-Whitney test. ***p ≤ 0.001, *p ≤ 0.05 n.s. not significant.

Next, we tested whether the levels of histone acetylation were affected in the absence of GATAD2B-NuRD complex. Using the PLA assay with antibodies against γH2AX and pan-acetyl H3 or Pan-acetyl H4, we found that depletion of GATAD2B or MBD3 and overexpression of RNaseH1 significantly increased levels of acetylated H3 or H4 upon DNA damage (Fig 4F and Fig EV10A). To validate these data further, we performed ChIP-qPCR analysis in DIvA U2OS cells and found that GATAD2B depletion led to increased levels of H4 acetylation around two DSBs (Fig 4G, H and Fig EV10B, C). Additionally, Western blot analysis showed that depletion of GATAD2B does not affect total level of H4 acetylation (Fig EV10D).

Overall, we conclude that GATAD2B-NuRD complex promotes histone deacetylation forming a boundary around DSBs.

### GATAD2B-NuRD complex boundary prevents chromatin hyper relaxation

NuRD complex has chromatin remodeling and deacetylation activity, which could promote chromatin condensation. Furthermore, transcription termination has been shown to be associated with repressive histone modifications and to recruit heterochromatin factors. To investigate whether R-loops and GATAD2B-NuRD complex and could affect chromatin structure upon DNA damage, we employed micrococcal nuclease (MNase) sensitivity assays. MNase is an endo-exonuclease that preferentially cleaves the accessible DNA linker between nucleosomes. MNase digestion of chromatin leads to a ladder of DNA fragments that can be visualized by agarose gel electrophoresis. Each band of the ladder corresponds to DNA protected by units of nucleosomes. The fastest migrating fragment (∼147 bp) represents DNA wrapped by a single nucleosome (mono-nucleosome), with fragments increasing by multiples of this length(Voong *et al*, 2017). Hence, more open and relaxed chromatin would result in higher number of mono-nucleosomes after MNase digest. First, we applied MNase digest to cells exposed to IR and observed that 10 min after IR, chromatin was more relaxed, as measured by increased number of mono-nucleosomes. Interestingly, RNaseH overexpression significantly increased MNase accessibility and digest upon DNA damage (Fig EV11A), suggesting that R-loop removal could lead to increased chromatin relaxation in early stage of DDR. Next, we performed MNase assay in cells depleted of GATAD2B and also detected significantly increased number of mono-nucleosomes, when compared to control cells upon DNA damage (Fig EV11B). To further validate the role of GATAD2B in chromatin remodeling upon DNA damage *in vivo*, U2OS cells were transfected with plasmids encoding for core histone H2B fused photoactivable GFP followed by laser micro irradiation(Smith *et al*, 2018). The thickness of GFP stripes was measured to indicate chromatin relaxation status. We found that depletion of GATAD2B led to significantly increased chromatin relaxation upon laser irradiation (Fig 5A-C), suggesting that GATAD2B-NuRD complex is required for chromatin condensation around DSBs.

**Fig 5.**
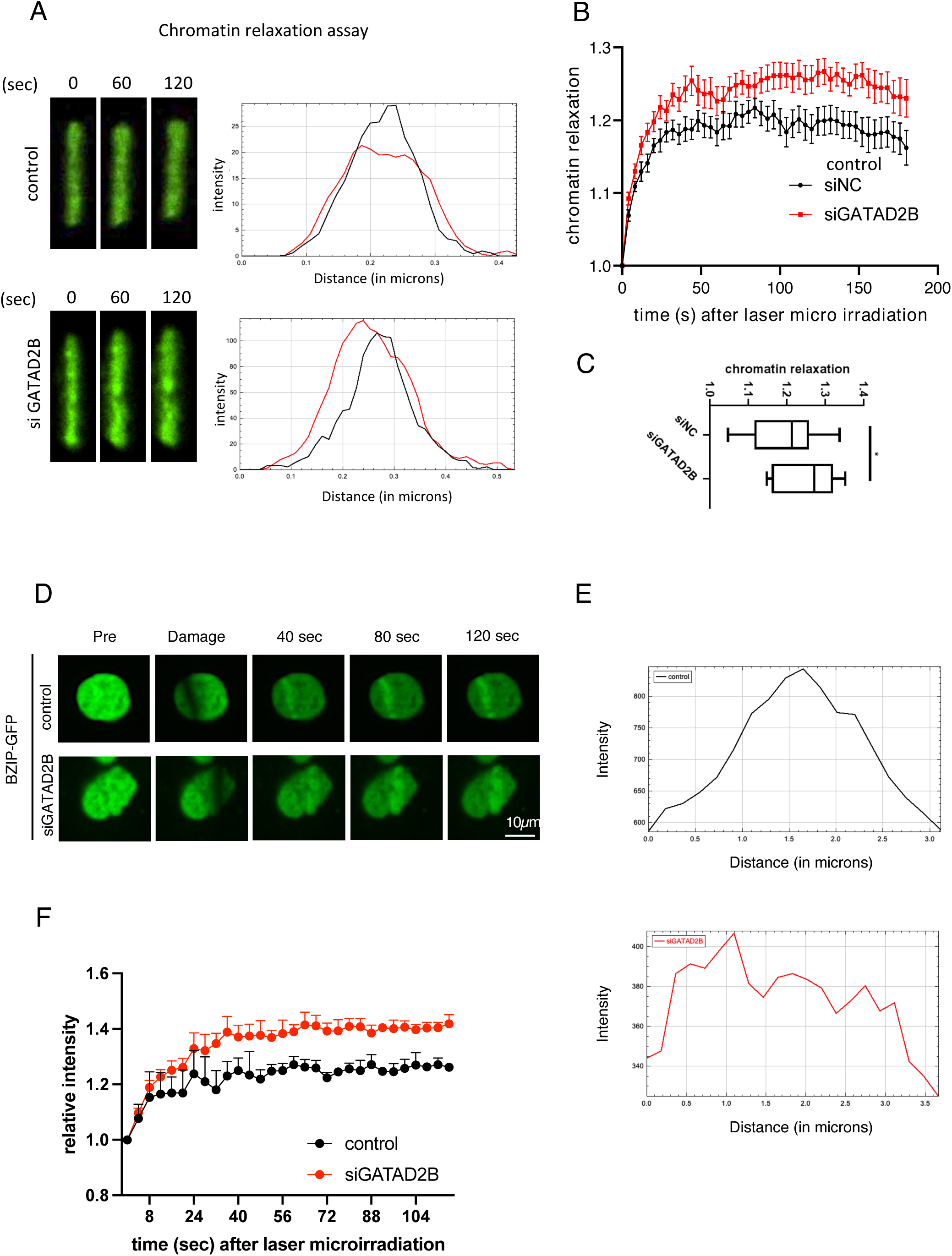
GATAD2B-NuRD complex facilitates chromatin condensation near DSBs. A) Left: Representative confocal images of the chromatin line that is simultaneously damaged by irradiation at 405 nm in U2OS wt or GATAD2B KD cells. Right: Intensity profiles showing thickness of the irradiated lines at 0 s (black) and 120 s (red) after damage induction. D) Graph showing kinetics of chromatin relaxation in wt U2OS and GATAD2B KD cells after DNA damage induction. C) Box plot showing the levels of chromatin relaxation at 120 s after laser irradiation in wt and GATAD2B KD cells. D) Representative confocal images showing recruitment of DNA-binding domain of BZIP from C/EBPa tagged with GFP to the sites of DNA damage induced by laser micro-irradiation at indicated time points in wt (control) and siGATAD2B cells. E) Representative plot profile showing BZIP-GFP signals across laser stripe, measured in micrones. F) Graph showing kinetics of BZIP-GFP recruitment to the sites of DNA damage induced by laser micro-irradiation at indicated time points in wt (control) and siGATAD2B cells.

Finally, in order to test chromatin status at DSBs, we employed BZIP DNA domain from C/EBPa transcription factor. C/EBPa has no known role in DDR and therefore its BZIP domain is not expected to be recruited to DSBs for the purpose of repair. Instead, it has been used previously, as a marker for open/relaxed/euchromatin(Smith *et al*, 2019). We have transiently transfected BZIP-GFP and employed laser micro-irradiation to monitor BZIP-GFP recruitment to DSBs in control and siGATAD2B cells. We show that depletion of GATAD2B results in more relaxed chromatin at DSBs as measured by increased intensity and wider BZIP-GFP stripes (Fig 5D-F).

Overall, our results suggest that GATAD2B-NuRD complex restricts chromatin relaxation around DSBs.

### GATAD2B-NuRD complex facilitates termination of DNA end resection and promotes HR

We show that the GATAD2B-NuRD complex is associated with HR prone DSBs. Initiation of HR depends on resection and production of ssDNA overhang, which is essential for D-loop formation and identification of homology regions in sister chromatid. Balanced levels of R-loops have been associated with correct end of resection in *fission yeast* (Ohle *et al*, 2016). We also show that GATAD2B-NuRD complex facilitates histone deacetylation and chromatin condensation. Hence, we tested whether the R-loops and GATAD2B-NuRD complex are also involved in end resection termination. Replication protein A (RPA) is ssDNA binding protein and the first responder to the presence of ssDNA overhang during resection. First, we measured the effect of transcription inhibition, RNAseH1 overexpression or GATAD2B depletion on the levels of phosphorylated RPA upon IR treatment. Notably transcription inhibition, overexpression of RNaseH1 or depletion of GATAD2B or MBD3 led to a significant increase in the number of pS4/S8 RPA32 foci (Fig 6A). In contrast, depletion of BRCA1 resulted in decreased number of pS4/S8 RPA32 foci (Fig 6A). Extended resection could be reflected also by increased size of pS4/S8 RPA32 foci. We employed FIJI software for the analysis of foci size and showed that transcription inhibition, overexpression of RNaseH1 or depletion of GATAD2B or MBD3 resulted in increased number of larger foci (as defined by prominence set to 2500 in Find Maxima tool) (Fig EV11C).

**Fig 6.**
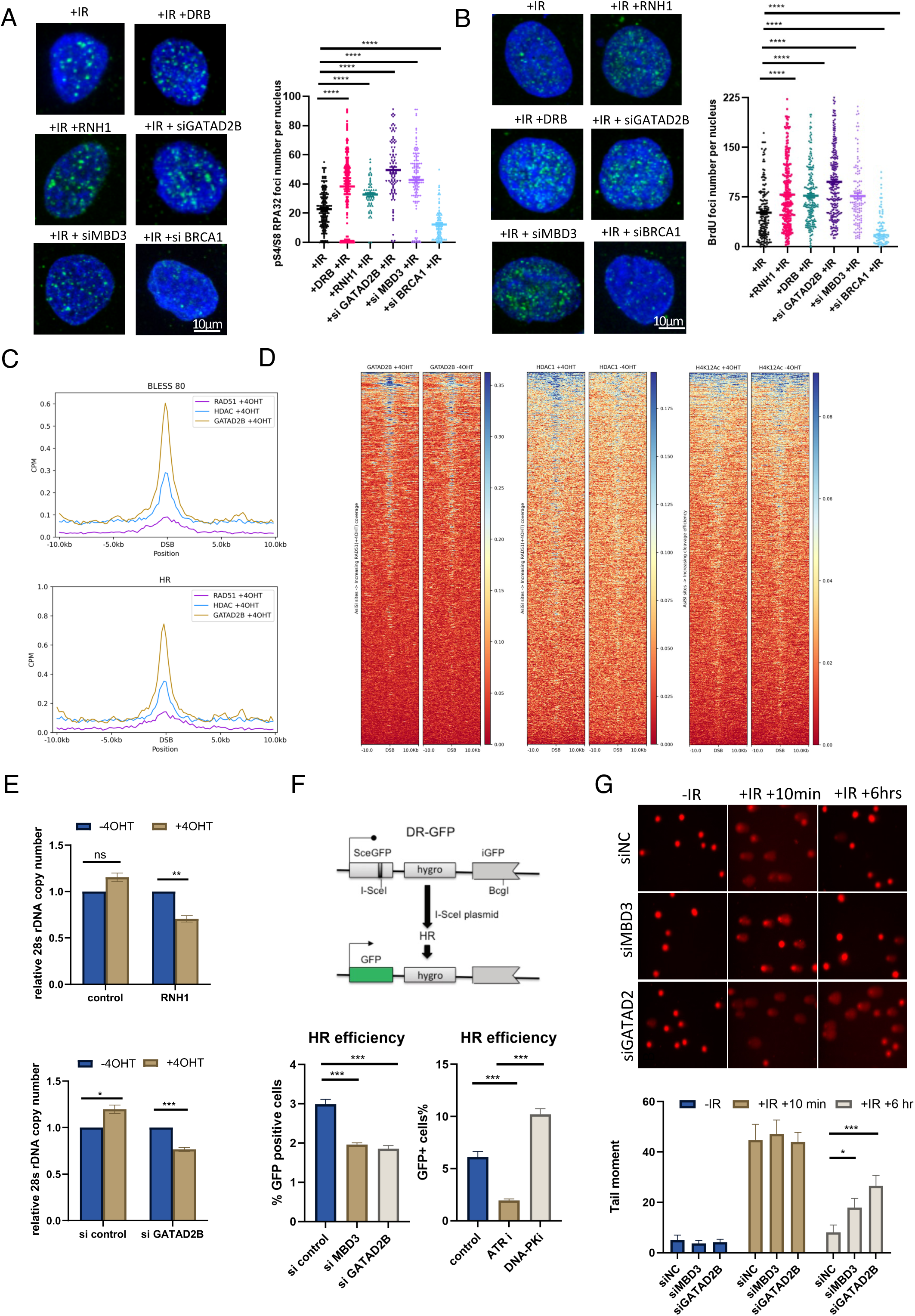
GATAD2B-NuRD complex facilitates termination of end resection and promotes HR. A) Left: representative confocal images showing immunofluorescence signals of phospho S4/S8 RPA32 in irradiated cells with overexpression of RNAseH1 or transcription inhibition (TLP3 or DRB) or depletion of GATAD2B and MBD3. Right: quantification of left, error bar = mean ± SEM, significance was determined using non-parametric Mann-Whitney test. ****p ≤ 0.0001 B) Left: representative confocal images showing immunofluorescence signals of BrdU in irradiated cells with transcription inhibition (TLP3 or DRB), overexpression of RNAseH1 or depletion of GATAD2B and MBD3. Right: quantification of left, error bar = mean ± SEM, significance was determined using non-parametric Mann-Whitney test. ***p ≤ 0.001, ** p ≤ 0.01, * p ≤ 0.05. C) Metagene profile showing ChIP-seq enrichment of GATAD2B, HDAC1 and RAD51 at cut AsiSI (as defined by BLESS technique) and HR prone cut AsiSI sites. D) Heatmaps showing GATAD2B, HDAC1 and H4K12Ac ChIP-seq and DRIP-seq read count over a 20 kb window centered on the DSB before (−4OHT) and after (+4OHT) DSB induction. DSBs are sorted according to decreasing coverage. See text for references. E) Bar charts showing change of 28s rDNA copy number upon 4OHT induction in cells expressing IPpOI, and overexpression of RNaseH (left) or knock down of GATAD2B (right) F) Top: Drawing indicating structure of HR reporter cassette. Bottom: bar charts showing HR repair efficiency under DSB inducted by I-SceI in U2OS cells containing HR reporter, and knock down of MBD3 and GATAD2B, inhibition of DNA-PK and ATR. G) Top: representative microscopy images showing comet assay of control (siNC), or GATAD2B and MBD3 knock down cells without IR, or irradiated with 5 Gy with 10 min recovery and 6 hr recovery. Bottom: quantification of top, error bar = mean ± SEM, significance was determined using non-parametric Mann-Whitney test. ***p ≤ 0.001, * p ≤ 0.05.

Intercalation of BrdU nucleotide into longer ssDNA can be visualized using anti-BrdU antibodies. We employed immunofluorescence assay and showed that transcription inhibition, overexpression of RNaseH1 or depletion of GATAD2B or MBD3 resulted in higher levels of BrdU signal, suggesting the presence of longer ssDNA. Depletion of BRCA1 resulted in decreased number of BrDU foci (Fig 6B).

Next, we have used available RAD51 ChIP-seq data from (Aymard *et al*., 2014) and overlapped it with our GATAD2B and HDAC1 ChIP-seq. Generated metagene profiles showed clear overlap between these three proteins around DSBs (Fig 6C and Fig EV12A-J). We have also generated a heatmap using AsiSI sites sorted based on RAD51 ChIP-seq coverage in 2.5kb flanking regions around DSBs and show that GATAD2B and HDAC1 ChIP-seq coverage is proportional to RAD51 coverage (Fig 6D).

Hyper-resection leads to longer ssDNA overhang, which could activate alternative DSB repair pathways, such as single-strand annealing (SSA) and micro-homology mediated end-joining (MMEJ) and could lead to a loss of genetic information(Zhao *et al*, 2020). Unrestricted DNA end resection could also result in increased ssDNA exposure, collapse of replication forks, deficiency of HR repair pathway and genome instability. Therefore, DNA end resection must be terminated at the point, when length of ssDNA overhang is sufficient to activate HR repair(Chen *et al*, 2013). To test whether GATAD2B-NuRD complex or R-loops could prevent activation of SSA or MMEJ pathways, we employed IPpOI-ER restriction enzyme digestion to induce site specific DSBs at 28s rDNA region(Berkovich *et al*, 2007), which consists of repetitive sequences. Activation of SSA or MMEJ repair pathways would result in loss of genetic information and reduced copy number of 28s rDNA. Indeed, RNaseH1 overexpression or GATAD2B depletion led to a reduction of 28s rDNA copy number upon DSBs induction by IPpoI (Fig 6E). Thus, our results indicate that GATAD2B-NuRD complex could facilitate termination of DNA end resection and consequently prevent loss of genetic information.

Next, we determined whether GATAD2B-NuRD complex is required for efficient DNA repair. U2OS DR-GFP and EJ5-GFP cells were employed as they possess HR and NHEJ reporter cassettes respectively (D’Alessandro *et al*., 2018). If DNA damage is induced by endonuclease cleavage; the repaired cells will generate GFP signal which can be measured and quantified by FACS. Notably depletion of GATAD2B and MBD3 decreased HR, but not NHEJ efficiency. As positive controls we used ATRi and DNA-PKi (Figs 6F and Fig EV13B). Furthermore, we used the HR reporter system in cells depleted of RNA helicases such as XRN2 and Senataxin, as well as RNAseH1 and show decreased efficiency of HR (Fig EV13A). These data suggest that the number of R-loops around DSBs has to be tightly regulated, which is in agreement with previous studies(Ohle *et al*., 2016). To extend these data further, we monitored the clearance of γH2AX in cells lacking GATAD2B and MBD3 or overexpressing RNAseH1 and found a delay in DNA repair as measured by increased levels of γH2AX (Fig EV13C and D). Finally, we employed comet assay to obtain a direct measurement of DNA repair. Again, depletion of GATAD2B and MBD3 resulted in significant accumulation of DNA tails corresponding to unrepaired DNA (Fig 6G).

Overall, our data show that GATAD2B-NuRD chromatin boundary prevents DNA end hyper-resection and promoting efficient DSB repair.

## Discussion

The process of DNA damage response involves the relaxation of chromatin, enabling the recruitment of DDR factors to the site of DNA lesions through DSB associated transcription. However, transcriptional repression and chromatin condensation also occur in the vicinity of DSBs. These observations imply the necessity for coordinated action between two distinct chromatin landscapes to ensure effective DSB repair. Notably, histone acetylation has emerged as a crucial regulatory mechanism governing chromatin structure upon DNA damage, with various histone acetyltransferases and deacetylases being observed at DSB sites.(Legube & Trouche, 2003). To achieve this coordination, the formation of a temporal boundary between open and condensed chromatin is likely critical. We show that R-loop formation around DSBs promotes the recruitment of the GATAD2B-NuRD complex to facilitate HDAC1-mediated histone deacetylation at DSBs at an early stage of DDR. This consequently creates a boundary between open and condensed chromatin, which in turn promotes termination of DNA end resection and efficient HR.

Chromatin remodeling is an important step in DDR. Relaxed chromatin is required for increased accessibility for DDR factors to broken DNA ends(Murr *et al*., 2006; Price & D’Andrea, 2013). It should be noted that heterochromatin markers, such as H3K9me3, were observed to be present next to DSBs, and several proteins involved in chromatin condensation, such as HP1 and G9a, were also found around DSBs(Ayrapetov *et al*., 2014; Kalousi *et al*., 2015; Yang *et al*., 2017). Interestingly, in non-damage condition, these two proteins are recruited to transcription termination loci to establish heterochromatin formation after R-loops dependent H3K9me2 deposition(Skourti-Stathaki *et al*., 2014). Moreover, R-loops play a role in regulating both chromatin relaxation and chromatin condensation. Additionally, a delicate balance of various proteins and histone modifications associated with heterochromatin and euchromatin has been observed at sequence-independent heterochromatin borders(Kimura & Horikoshi, 2004; Wang *et al*, 2014). Therefore, it is plausible that R-loops formed at DSBs act as boundary elements, facilitating the recruitment of the GATAD2B-NuRD complex to promote histone deacetylation. Potentially, this deacetylation could allow for the availability of lysine residues on histones for the deposition of other modifications, such as methylation by G9a. Consequently, this could facilitate the loading of HP1, which in turn recruits SUV39H to methylate neighboring histones and establish and propagate heterochromatin structure.

Interestingly, occupancy of GATAD2B and HDAC1, whose localization to DSB is dependent on R-loops, shows negative correlation with H4K12ac around DSBs(Clouaire *et al*., 2018a). This finding suggests that acetylated H4K12 may be a potential substrate of the NuRD complex around DSBs. Notably, H4K12Ac has been found to be associated with transcriptionally active regions(LeRoy *et al*, 2008). Additionally, RNAPII pausing has been detected around DSBs(Pankotai *et al*, 2012), and *de novo* transcription at these loci also needs to be terminated. Thus, we hypothesized that the HDAC1-containing NuRD complex may promote deacetylation H4K12 to facilitate DSB associated transcription termination.

The NuRD complex is a chromatin-remodeling complex that consists of several subunits, including GATAD2A and GATAD2B. These subunits define specific NuRD subcomplexes that play a crucial role in regulating the chromatin environment surrounding DSBs(Spruijt *et al*., 2016). GATAD2A-NuRD is localized at DSBs in a ZMYND8-depedent manner, which is a histone acetylation reader and knockdown of ZMYND8 impairs HR efficiency(Gong *et al*., 2017; Spruijt *et al*., 2016).

Another component of the NuRD complex, CHD4, has been found in close proximity to DSBs(Gong *et al*., 2015). Interestingly, CHD4 promotes chromatin relaxation upon DNA damage, either in a NuRD-dependent or independent manner(Hou *et al*, 2020; Smith *et al*., 2018). It is plausible that CHD4-dependent local chromatin relaxation is necessary for *de novo* transcription and R-loop formation around DSBs. By promoting local chromatin relaxation, CHD4 plays a vital role in facilitating the formation of an appropriate chromatin landscape necessary for transcriptional activation, R-loop formation, and the recruitment of GATAD2B-NuRD, which in turn promotes chromatin condensation.

One crucial aspect of homologous recombination (HR) is DNA end-resection, which is facilitated by an open chromatin landscape(Costelloe *et al*, 2012; Pai *et al*, 2014). However, excessive end resection can result in long single-stranded DNA (ssDNA) overhangs triggering the activation of single-strand annealing (SSA) and microhomology-mediated end joining (MMEJ) pathways(Zhao *et al*., 2020). These pathways can lead to significant loss of DNA, particularly in repetitive regions. Therefore, it is crucial to restrict end resection to approximately 2 kilobases (kb) from DSBs to ensure accurate HR repair(Cohen *et al*., 2018; Mimitou *et al*, 2017). Our data suggest that the formation of a chromatin boundary dependent on the GATAD2B-NuRD complex could play a role in maintenance of the appropriate range of end resection, preventing excessive loss of genetic material and ensuring accurate and effective DNA repair.

Overall, our study provides evidence for the establishment of the temporal boundary between open and condensed chromatin in R-loop and GATAD2B-NuRD complex-dependent manner (Fig 7). These findings have important implications for understanding the complex mechanisms underlying the regulation of DDR and could pave the way for the development of novel therapeutic strategies for various genetic disorders.

**Fig 7.**
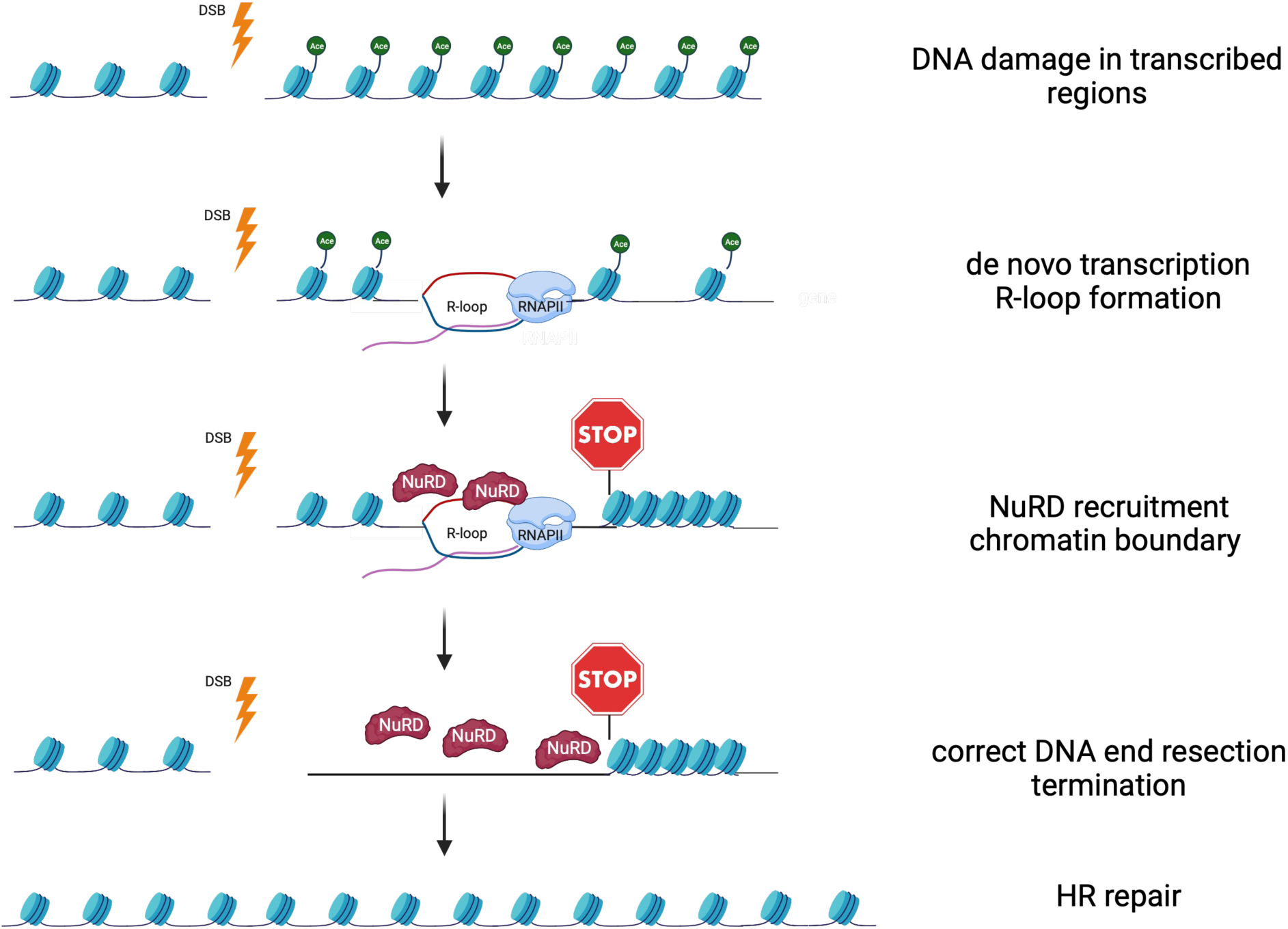
The model of chromatin boundary around DSBs established by GATAD2B-NuRD complex and R-loops. GATAD2B-NuRD complex associates with DSBs in a transcription and R-loop-dependent manner, creating a boundary and facilitating histone deacetylation and chromatin condensation around DSBs. Its presence near damaged loci is required for controlled end resection termination. The lack of the GATAD2B-NuRD complex leads to chromatin hyper-relaxation and extended DNA end resection, causing failure in HR repair.

## Methods

### Tissue Culture

All human cell lines applied in this study (HEK293T, HeLa, U2OS, DR-U2OS, EJ5-U2OS, AsiSI-ER) were cultured in DMEM media with 10%-20% fetal bovine serum, L-glutamine and penicillin/streptomycin. All human tissues were cultured under the condition 37℃, 5% CO2.

### S9.6 Immunoprecipitation

The protocol for this assay was described previously. Briefly, nuclei from ionized radiated or non-treated HEK293T cells were extracted by cell lysis buffer (5mM PIPES pH8.0, 0.5% NP-40 and 85mM KCl) on ice for 10 min. Isolated nuclei were then resuspended in nuclear lysis buffer (10 mM Tris-HCl pH 7.5, 200 mM NaCl, 2.5 mM MgCl2, 0.2% sodium deoxycholate [NaDOC], 0.1% SDS, 0.05% sodium lauroyl sarcosinate [Na sarkosyl] and 0.5% Triton X-100), followed by 10 min sonication (Diagenode Bioruptor). The extracts were then precleared, after which S9.6 antibodies were added for immunoprecipitation. Protein G dynabeads were then applied to bind with antibody, followed by 5 times washing and elution.

### Treatment of inhibitors, transfection of plasmids and siRNA

When indicated, cells were treated with 1 μM triptolide or 100 μM 5,6-dichloro-1-beta-D-ribofuranosylbenzimidazole (DRB) to inhibit transcription 2 hours before IR treatment or other assays. In the RNAse H overexpression group, cells were transfected with plasmids encoding RNAse H1 using Lipofectamine 3000 48 hours prior to the subsequent step. For siRNA transfection, siRNA was delivered into cells via Lipofectamine RNAiMAX 48 hours before IR treatment or other specified assays. For inhibitors, Cells were treated AZD 5305 (10 nM, 6 hrs) to inhibit PARP1, Ku-55933 (10μM 2 hrs) to inhibit ATM and AZ20 (500 nM 2 hrs) to inhibit ATR.

### Proximity Ligation Assay (PLA)

These experiments were performed according to manufacturers’ instructions. Specifically, cells were fixed by 4% paraformaldehyde solution at room temperature for 10 min, followed by PBS washing and 10 min lysis by 0.2% Triton-X 100.Cells were then blocked in PLA blocking buffer at 37℃ for 1 hr, after which primary antibodies were diluted and incubated with samples at cold room overnight. Those primary antibodies were then captured by PLA probes which were then ligated, followed by amplification by polymerase. For S9.6 related PLA, cells were pre extracted by CSK buffers (10 mM PIPES, 100 mM NaCl, 300 mM Sucrose,3 mM MgCl2, 1 mM EGTA, 0.7% Triton X-100, 1×protease inhibitor cocktail) at cold room for 15 min prior to fixation according to a previous report, while 1 hr incubation at room temperature with RNase treatment solution (0.1% BSA, 3 mM MgCl2, 1:200(v/v) RNAse T1, 1:200 (v/v) ShortCut RNAse III in PBS) was required for each sample after fixation.

### Laser micro-irradiation

Cells were transfected with related GFP or mCherry-tagged plasmids 48 hrs before experiments, and then seeded on glass bottomed dish (NEST) and pre-treated with 10 μM BrDU 24 hrs prior to irradiation. UV laser (16 Hz pulse, 40% laser output) generated from the Micropoint System (Andor) was then applied to induce DNA damage. Live-cell images were then taken by a Nikon A1 confocal imaging systems for every 5 or 10 seconds. Images were then analyzed by ImageJ and intensities of damage, non-damage and background areas were measured respectively to calculate relative intensities of stripped areas.

### Chromatin Immunoprecipitation (ChIP)

AsiSI-ER U2OS cells were treated 400 nM 4-Hydroxytamoxifen (4-OHT) 4 hrs before 10 min fixation by 1% formaldehyde solution. Fixed cells were then quenched by 125mM glycine at room temperature for 10 mins and scrapped at cold room. Later, tissues were lysed in cell lysis buffer mentioned above and nuclei lysis buffer (50 mM Tris-HCl pH8.0, 1% SDS and 10 mM EDTA) respectively on ice and sonicated for 15min. Samples were then spun for 10 mins at 13000 rpm to remove cell debris. The supernatant was the pre-cleared by protein G dynabeads and diluted by dilution buffer (16.7 mM Tris-HCl pH8.0, 0.01% SDS, 1% Triton-X 100, 167mM NaCl, 1mM EDTA) in a 1:4 ratio. Diluted samples were then aliqoted into several ChIP samples. Antibodies and beads were then added for immunoprecipitation, and beads were washed 5 times before elution in ChIP elution buffer (1% SDS and 100 mM NaHCO3) at room temperature for 30 min. Eluted samples were then de-crosslinked at 65℃ overnight and immunoprecipitated DNAs were then extracted by using Phenol-Chloroform extraction methods for following qPCR and sequencing experiments. For primers see Table S1.

### MNase Assay

Cells were collected in clean Eppendorf tubes, lysed in buffer A (50 mM Tris-HCl pH 8.0, 0.1% Triton-X 100, 150 mM NaCl, 1mM EDTA) to extract chromatin. Chromatin pellets were then resuspended with digestion buffer (15 mM Tris-HCl pH 7.5, 60 mM KCl, 15 mM NaCl, 0.25 M Sucrose, 1mM CaCl2, 0.5 mM DTT) with 100 units of MNase (NEB), incubated at 37℃ for 5 min. Digested DNA was then purified by Phenol-Chloroform extraction methods and separated by 1.2% agarose gel electrophoresis. Each representative image was selected from three biological repeats.

### Chromatin relaxation and DNA accessibility assay

The chromatin relaxation assay was performed as in(Smith *et al*, 2023). This assay uses PAGFP tagged histone H2B to mark a chromatin region by local photoactivation. This happens at the same time as laser irradiation at 405 nm causes DNA damage. The assay measures the thickness of the photoactivated line to estimate how much the chromatin condenses at the damage sites.

BZIP DNA domain from C/EBPa transcription factor was fused to GFP(Smith *et al*., 2023). Cells were transiently transfected and subjected to laser micro-irradiation and monitored BZIP-GFP recruitment to DSBs in control and siGATAD2B cells.

### HR and NHEJ Reporter Assay

DR-U2OS or EJ5-U2OS reporter cell lines were transfected siRNA by Lipofectamine RNAi Max (Life Technology) 48 hrs prior to I-SceI lentivirus transduction. After infection by lentivirus containing I-SceI, reporter cell lines were cultured at 37 ℃ for 48 hr before trypsinized and resuspended by PBS. Samples were then analyzed by a BD Accuri C6 flow cytometer.

### Immunofluorescence

Cells were seeded on glass bottomed dish or coverslips 24 hr prior to ionic irradiation, followed by CSK pre-extraction and fixation which were mentioned above. Coverslips were then blocked with 3% BSA at 37 ℃ for 1 hr, after which samples would be incubated with primary antibodies overnight at cold room and secondary antibodies at 37 ℃ for 1 hr subsequently. For BrdU IF, cells were incubated with 20 μM BrdU 24 hr prior to 5 Gy IR treatment, followed by 2 hrs recovery before the CSK extraction step. Cells were mounted on slides with Fluoroshield Mounting Medium with DAPI (Abcam). Images were then acquired by an Olympus Fluoview FV1200 confocal microscope.

### Comet assay

The assay was performed according to manufacturers’ instruction (bio-techne). Briefly, 5*10^3^ cells in 5 μl PBS were mixed with with 50 μl molten LMAgarose at 37°C, and spread onto CometSlide. CometSlides were then transferred to 4°C, before lysed in lysis buffer (2.5 M NaCl, 100 mM Na2EDTA, 10 mM Tris, with final pH 10.0) with 10% DMSO and 1% Triton-X 100 at 4°C for at least 2 hrs. Samples were then immersed into electrophoresis buffer (300 mM NaOH, 1 mM EDTA) to unwind for 30 min. The electrophoresis was performed under 21V, 200 mA condition at cold room. Slides were then washed 4 times and stained by Propidium Iodide for 30 min before imaging on a fluorescence microscope.

**For all reagents used in this study see Supplementary Table.**

### Bioinformatic analyses

#### ChIP-Seq data processing

GATAD2B +4OHT, GATAD2B -4OHT, HDAC1 +4OHT, HDAC -4OHT samples were sequenced using Illumina NextSeq 500 (single end 151 bp reads). Read quality of the fastq files was checked using FastQC (https://www.bioinformatics.babraham.ac.uk/projects/fastqc/) before and after adapter trimming. The trimmed reads were then mapped to hg19 genome following the standard ChIP seq-pipeline. The classic ChIP-seq pipeline consists of BWA (http://bio-bwa.sourceforge.net/) for alignment and samtools (http://www.htslib.org/) for duplicate removal (rmdup), sorting (sort) and indexing (index). deepTools (https://deeptools.readthedocs.io/en/develop/)(Ramirez *et al*, 2016) bamCompare was used to calculate log2 fold change at each nucleotide position between +4OHT and -4OHT for GATAD2B and HDAC1 samples. For peak calling, we have first extracted peaks with Input as control using MACS2 software. We then compared the replicates using ChIP QC software. PCA plots comparing coverage across consensus peaks show that at least two replicates per condition cluster together. For the downstream analysis we used one of the clustering replicates for each condition.

Read count normalization was applied as part of bamCompare to normalize for sequencing depth between samples. H4K12Ac +4OHT and H4K12Ac -4OHT ChIP-seq samples from (Clouaire *et al*, 2018b) ( E-MTAB-5817) was processed using the same approach for downstream analysis plots.

RNA Pol II S5P ChIP from a previous study(Cohen *et al*., 2018) (E-MTAB-6318) was processed using the classical ChIP pipeline as well. Read count coverage was calculated for each the 80 inducible DSB site (+-5kb) using bedtools multicov (https://bedtools.readthedocs.io/en/latest/)(Quinlan & Hall, 2010). The top 20 sites with highest coverage were annotated as high transcriptional activity and the bottom 20 as low transcriptional activity.

#### BLESS Seq data processing

BLESS Seq (E-MTAB-5817) was processed using the same protocol as detailed in previous publication(Clouaire *et al*., 2018a). Read count coverage was calculated for all annotated DSBs (+-500bp) using bedtools multicov. The sites were then ordered based on read count coverage for representing cleavage efficiency of DSB sites.

#### DRIP Seq data processing

DRIP Seq samples +/- 4OHT from a previous research work(Cohen *et al*., 2018) (E-MTAB-6318) was processed using the classical ChIP pipeline. Log2 Fold Change was calculated using bamCompare at each nucleotide position with readcount normalization. bamCompare outputs were exported as bigwig files for further downstream analysis.

#### Metagene Profiles

computeMatrix operation of deepTools was used to calculate the average profile of log2Fold Change (+4OHT/-4OHT) from the bigwig files generated using bamCompare. The bin size was set to 200 and region length was set to +/-10kb with reference to DSB. The average profile matrices created for GATAD2B, HDAC1 and DRIP-Seq were then combined and plotted using matplotlib package of python.

#### BoxPlots

Read count coverage around DSBs (+/- 2.5 kb) was calculated using bedtools multicov and then normalized to CPM using total read count of respective samples. The plots were made with matplotlib package of python. Wilcoxon two sample test was used to determine statistical significance with p-value cut-off set at 0.01. Statistical tests were run using scipy python package.

#### IGV profiles and Heatmaps

Coverage files containing CPM normalized read count per nucleotide position was generated for each sample using deeptools bamCoverage. The coverage files were exported in bigwig format and loaded into IGV browser (https://software.broadinstitute.org/software/igv/) for visualizing the normalized read density around break sites.

The coverage files were also used to make the heatmaps comparing cleavage efficiency and read density of HDAC1, GATAD2B, R-loops and H4K12Ac around DSB in 4OHT treated samples. plotHeatmap function of deepTools was used to make the heatmaps with regions set as all annotated DSBs arranged in ascending order of cleavage efficiency.

## QUANTIFICATION AND STATISTICAL ANALYSIS

For IF and PLA, at least fifty cells were analysed. Foci numbers were quantified by cell profiler software. For laser, fluorescence intensity was measured by software imageJ. Unless indicated, two-tailed Student’s t tests were applied for statistical analysis, while * is refers to p < 0.05, ** is refers to p < 0.01, *** is refers to p < 0.001 and **** is refers to p < 0.0001.

## Acknowledgements

We would like to thank all the members of the Gullerova and Zhu labs for their help and advice throughout this study. We grateful to Dr Rebecca Smith for her help with chromatin relaxation assays. We also thank to Dr Jun Zhang for his help with microscopy imaging and stimulating scientific discussion and Xingkai He with her assistance with sample preparation for qRT-PCRs. We are grateful to Dr Gaelle Legube for providing HR and NHEJ prone AsiSI cut site genome coordinates.

## Author Contributions

Z.L. designed and performed most of the experiments, with the assistance of Y.W. K.A. performed all of the bioinformatic analysis. W.Z. advised and supported part of the study. M.G. designed and supervised the project wrote and edited the manuscript. All the authors reviewed and approved the final version of the manuscript.

## Data Availability

All data are available in the paper and the Supplementary Data.

"The mass spectrometry proteomics data have been deposited to the ProteomeXchange Consortium via the PRIDE partner repository with the dataset identifier PXD042271". Data can be accessed via reviewer account details: Username: Password: r9fVIgEC

ChIP-seq datasets are deposited in GEO database under accession number GSE230710 and accessible with token yjyzssmsjjydxiv.

---

**Expanded View Figure 1.**
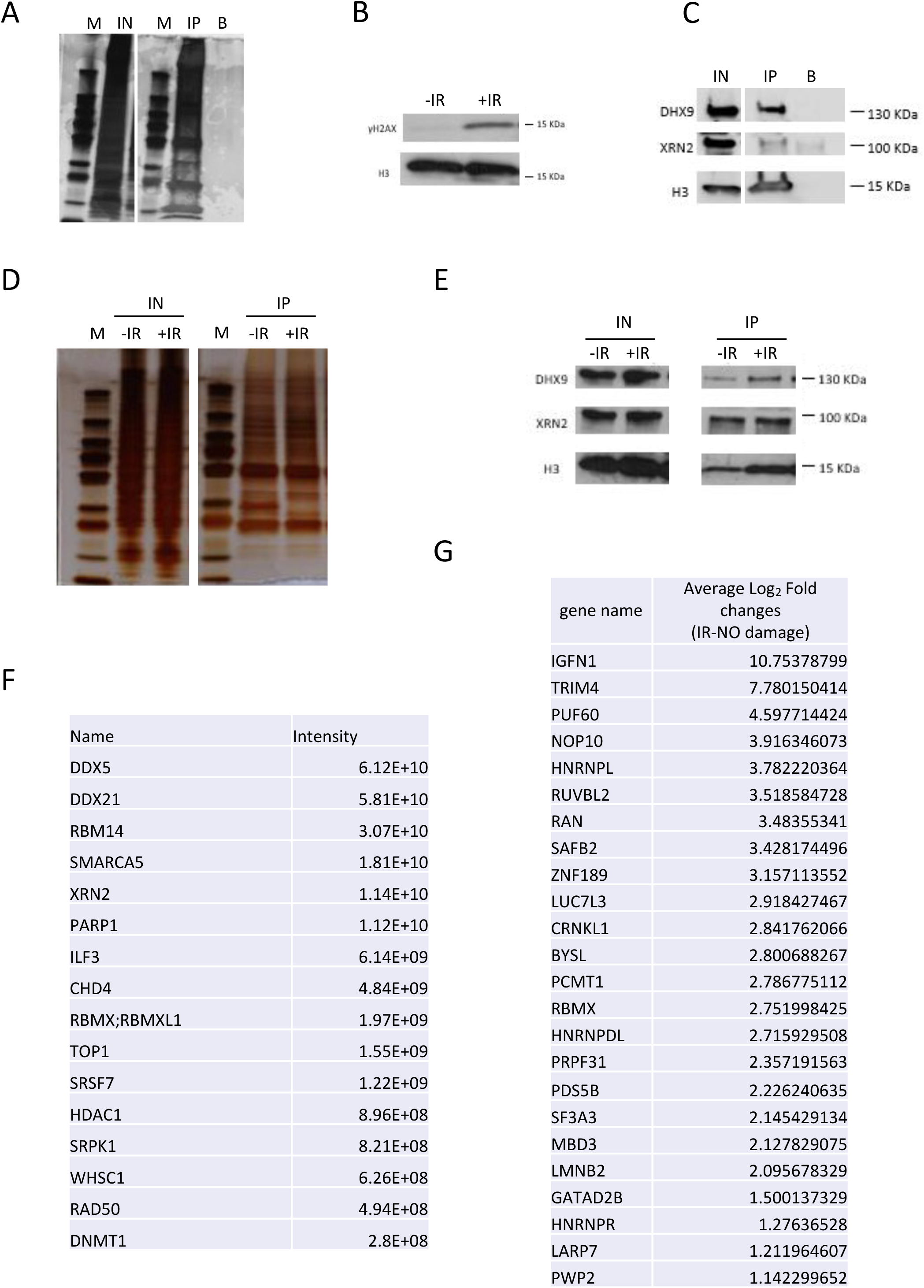

**Expanded View Figure 2.**
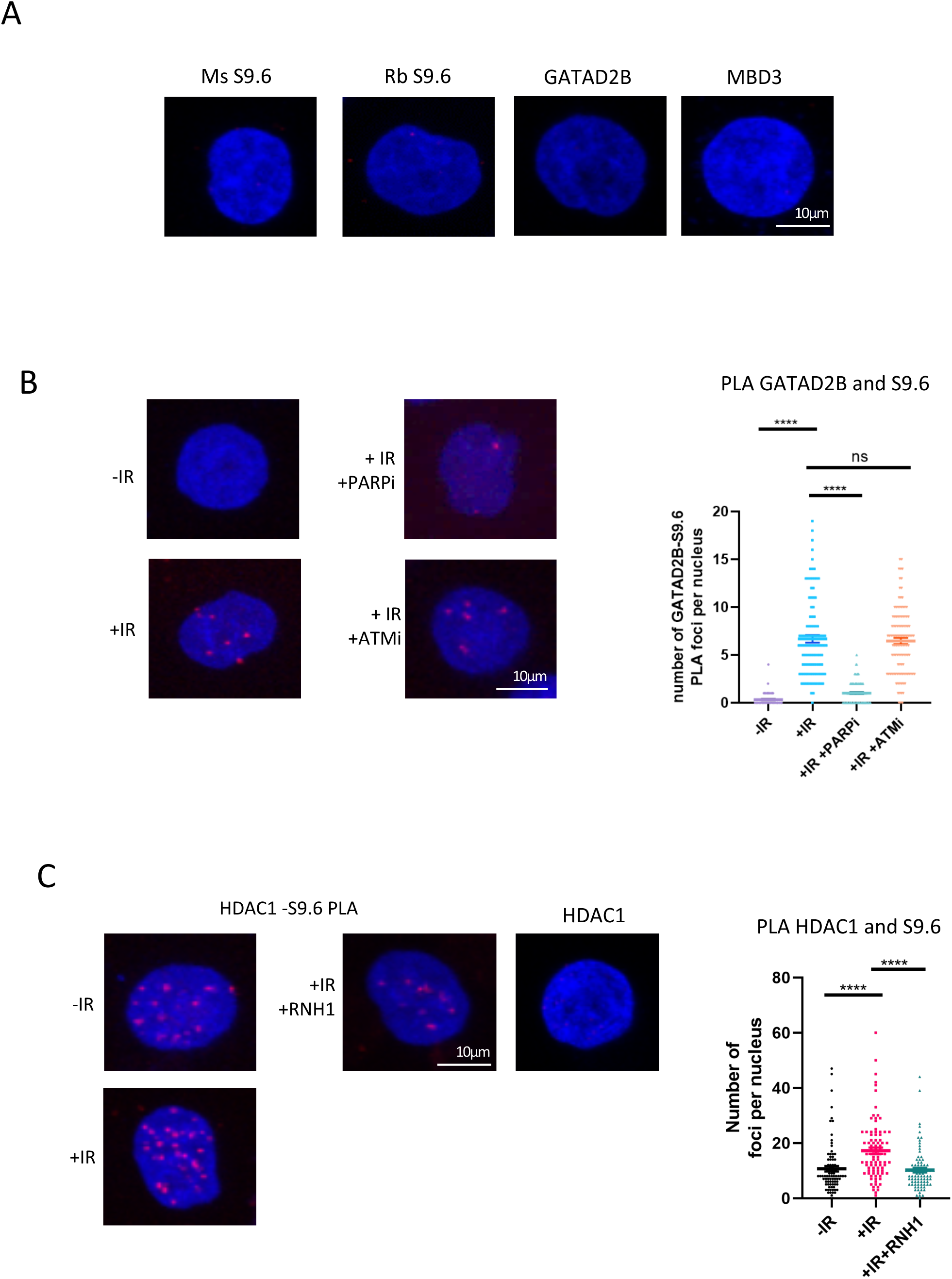

**Expanded View Figure 3.**
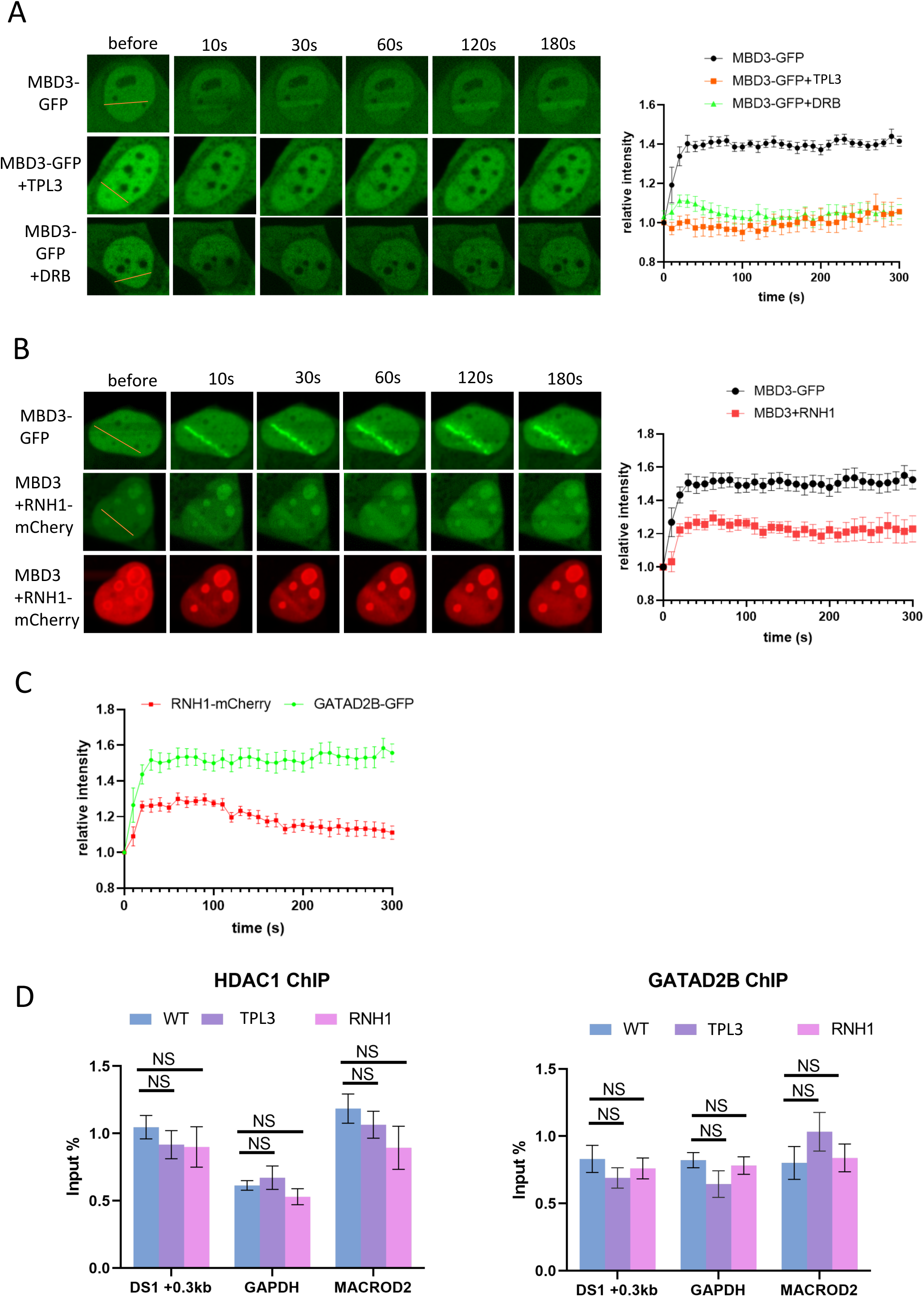

**Expanded View Figure 4.**
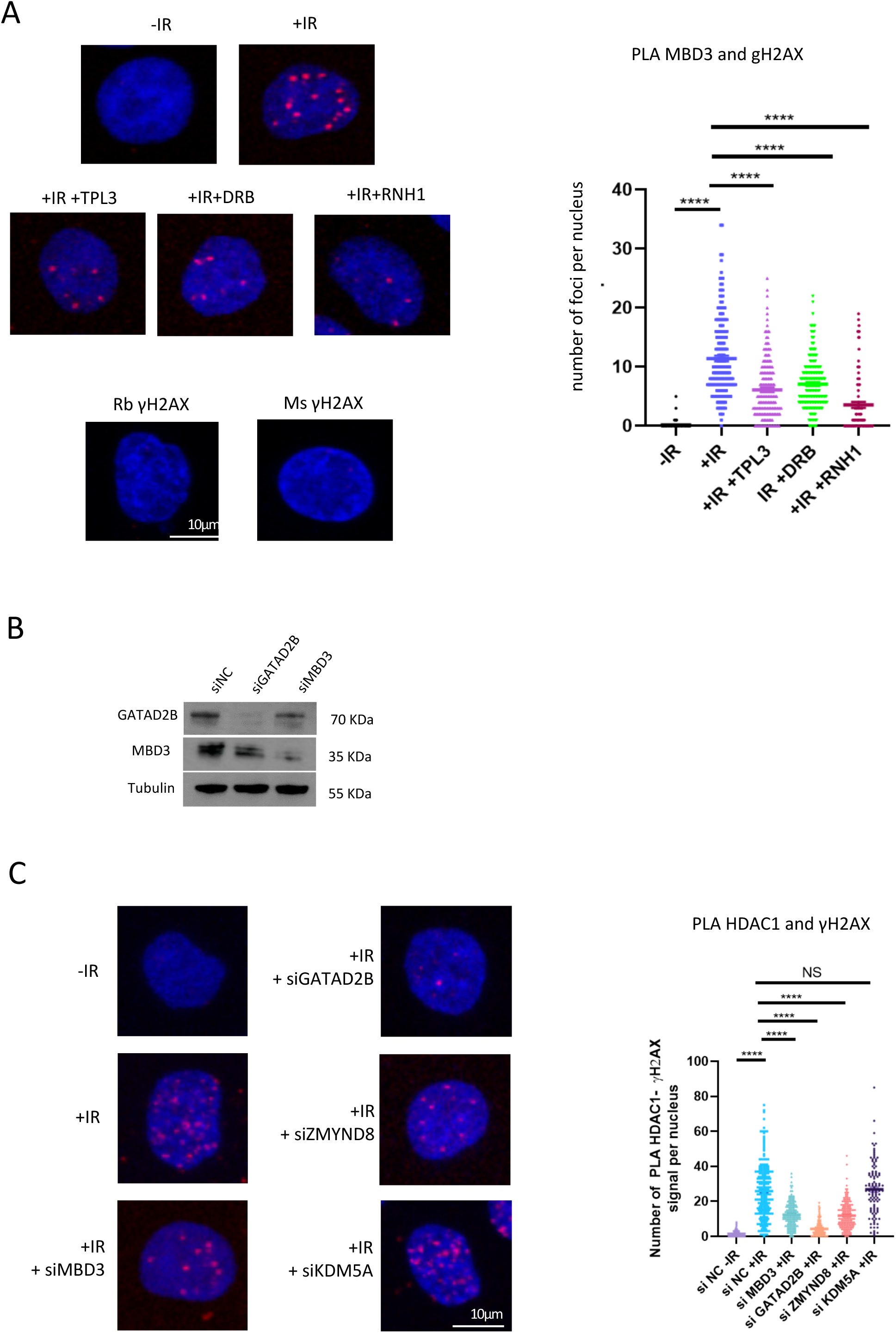

**Expanded View Figure 5.**
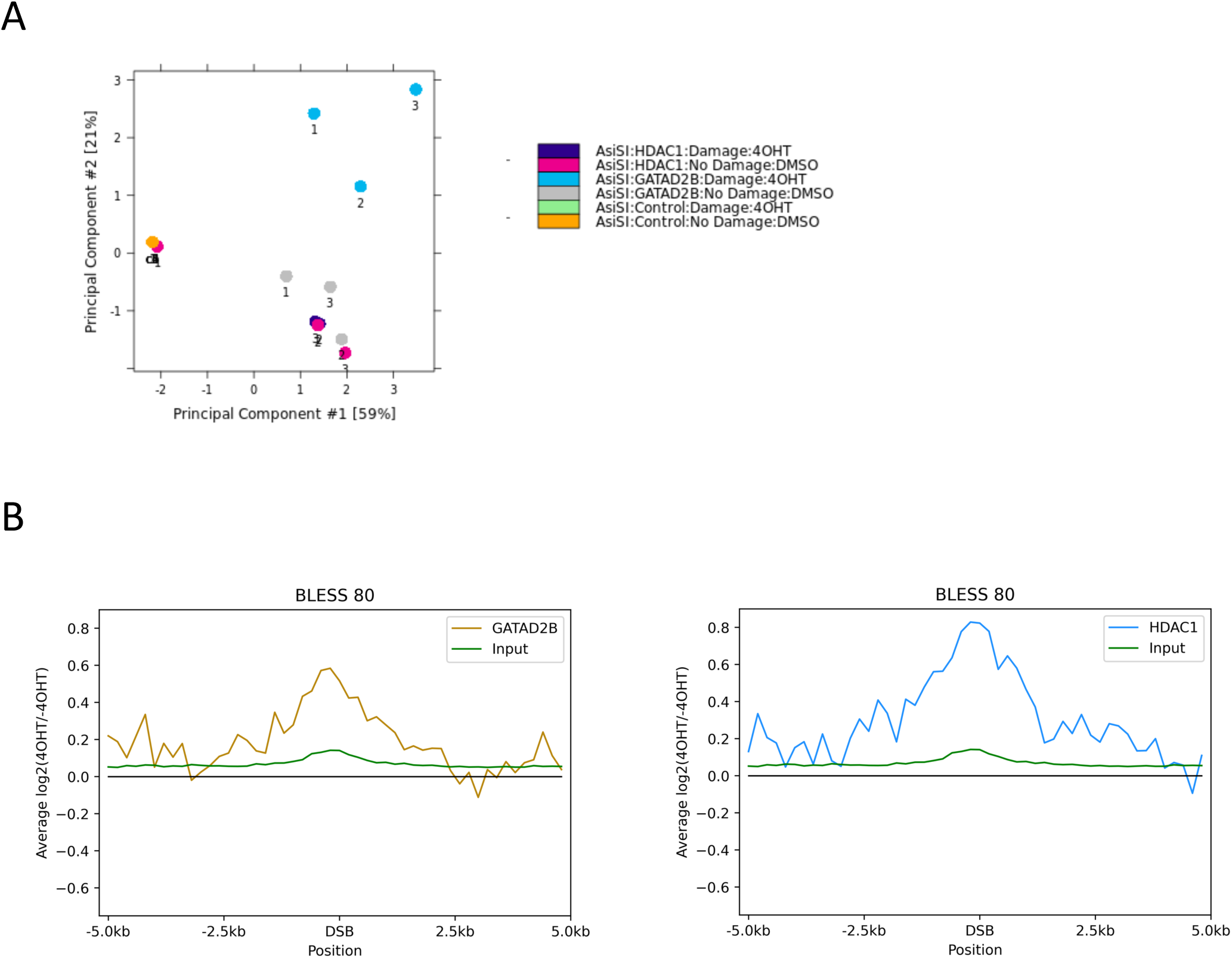

**Expanded View Figure 6.**
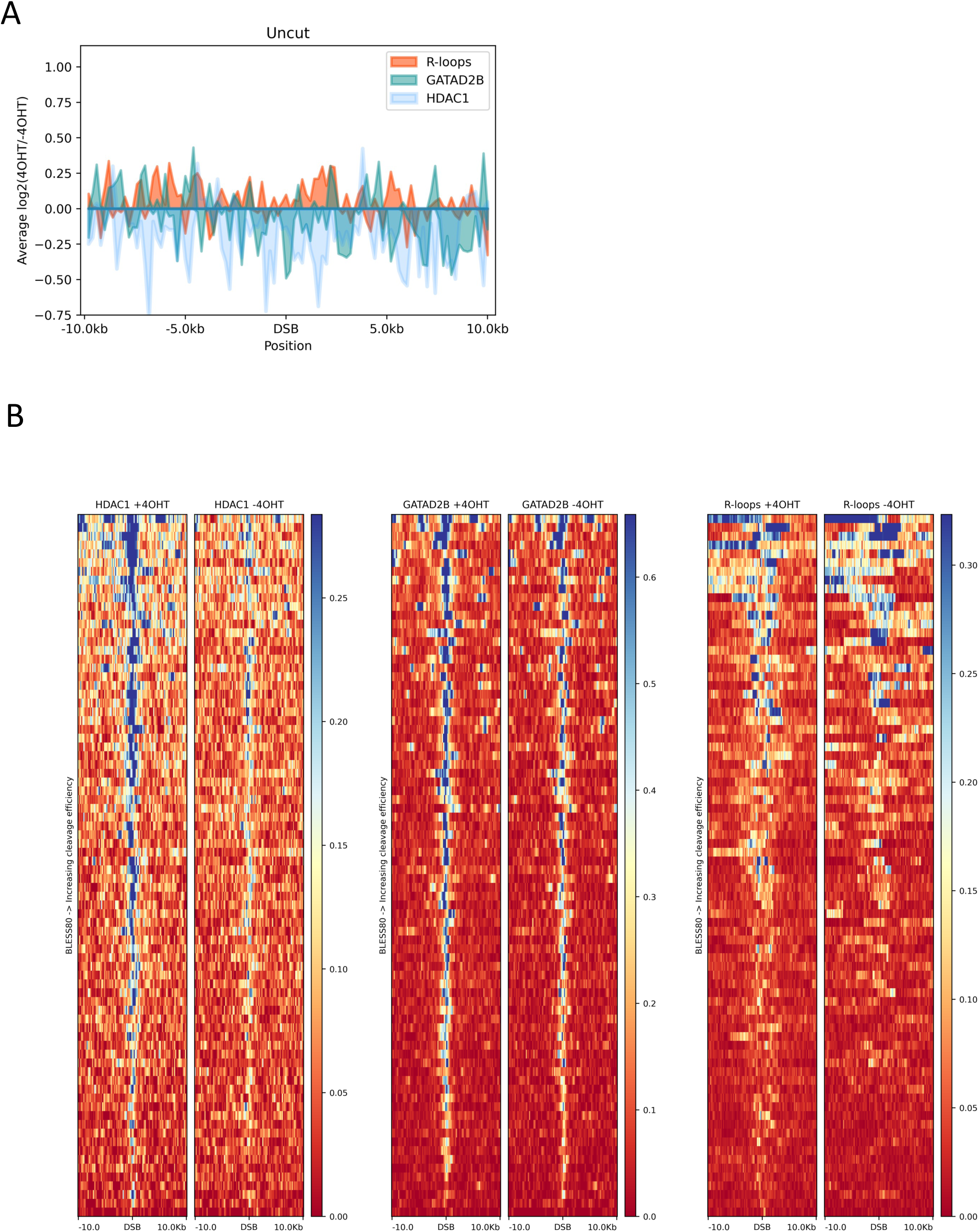

**Expanded View Figure 7.**
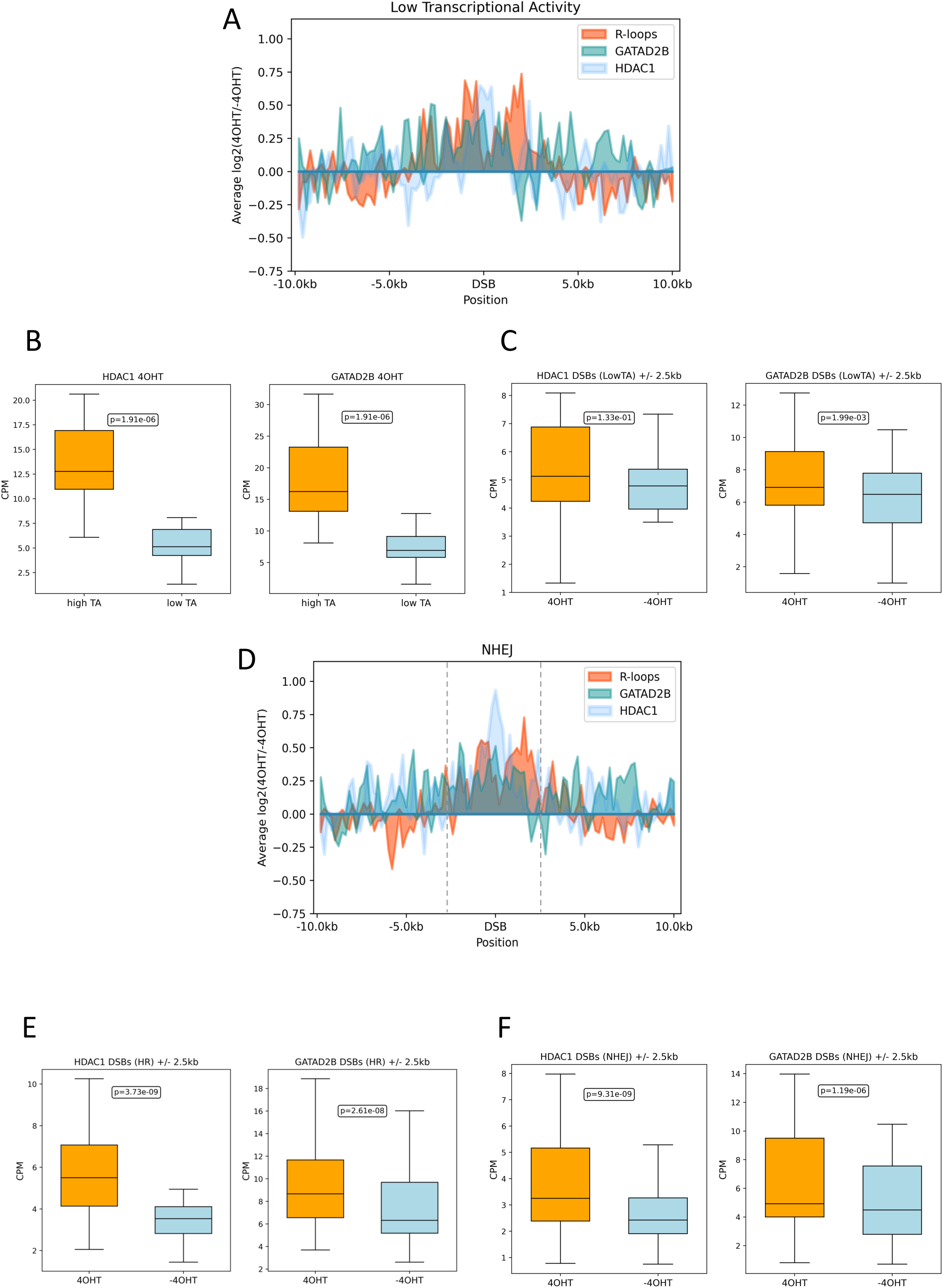

**Expanded View Figure 8.**
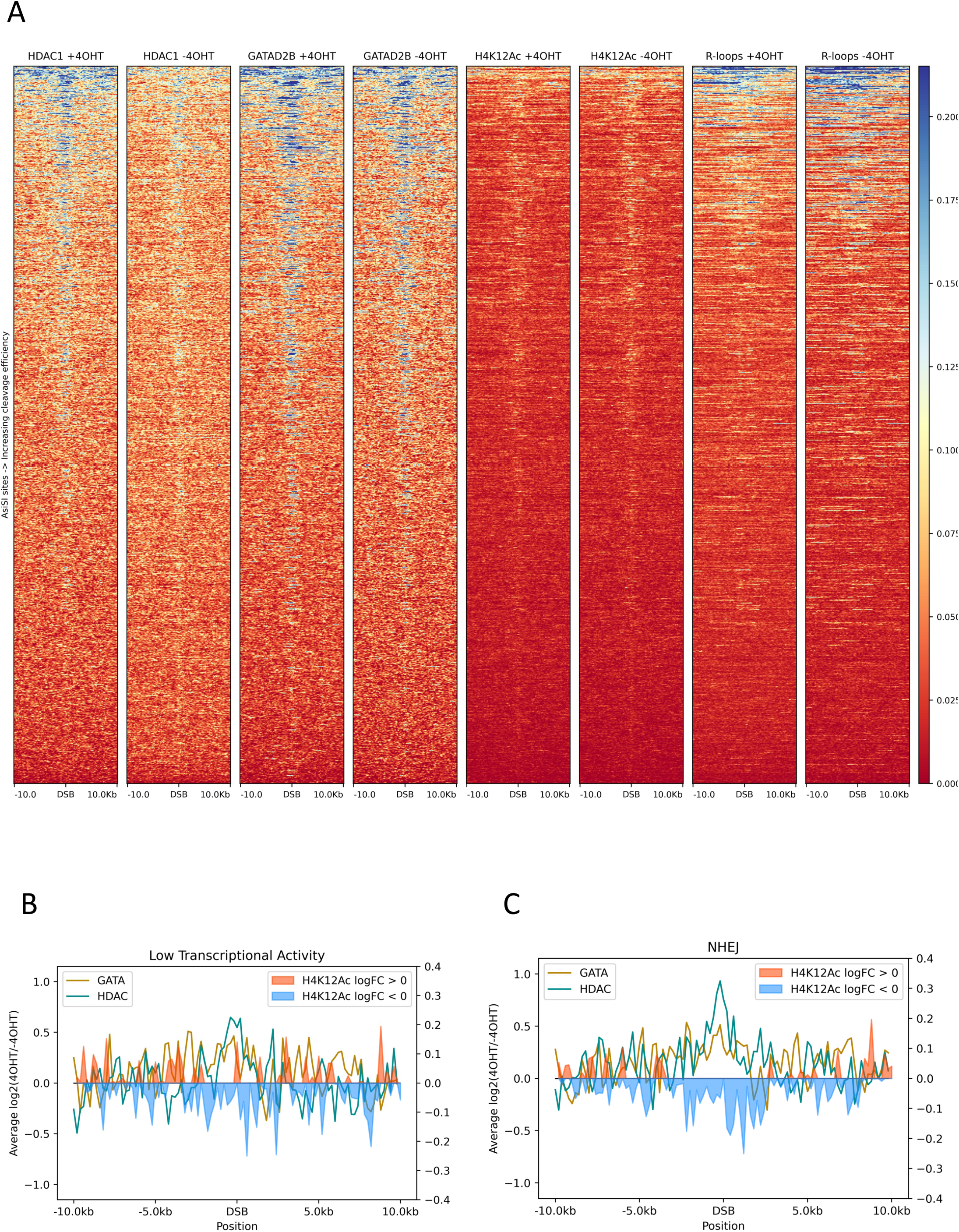

**Expanded View Figure 9.**
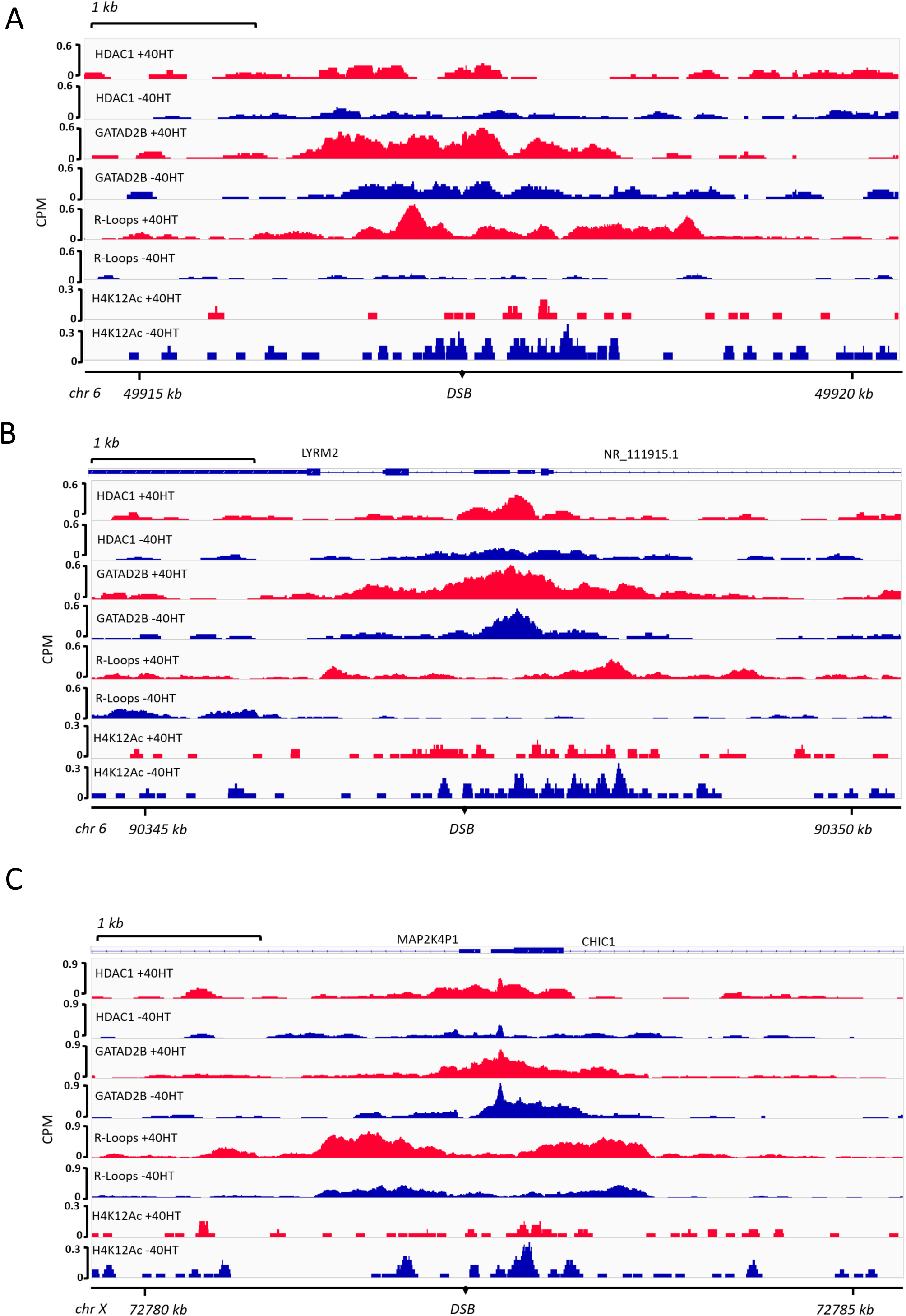

**Expanded View Figure 10.**
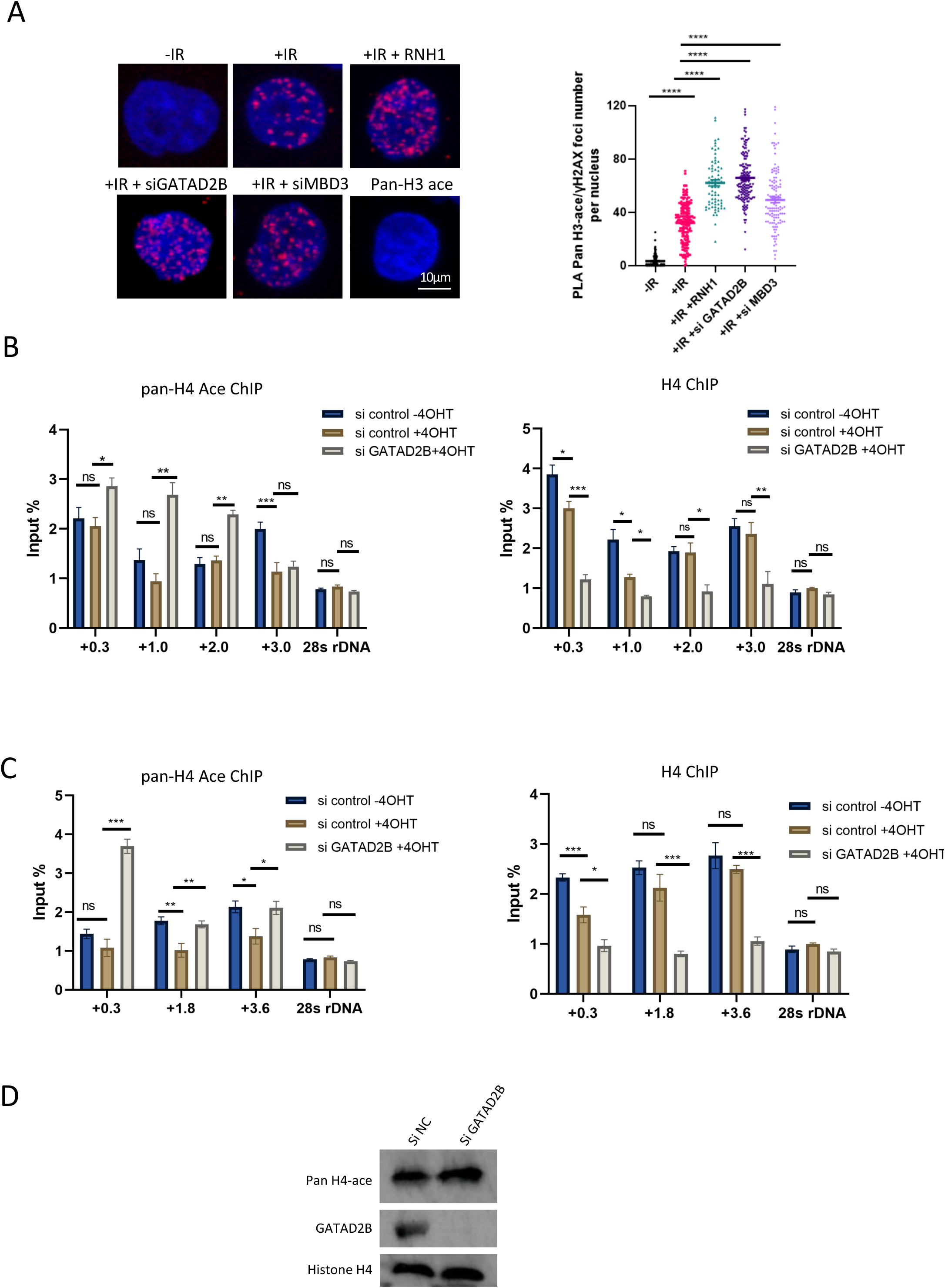

**Expanded View Figure 11.**
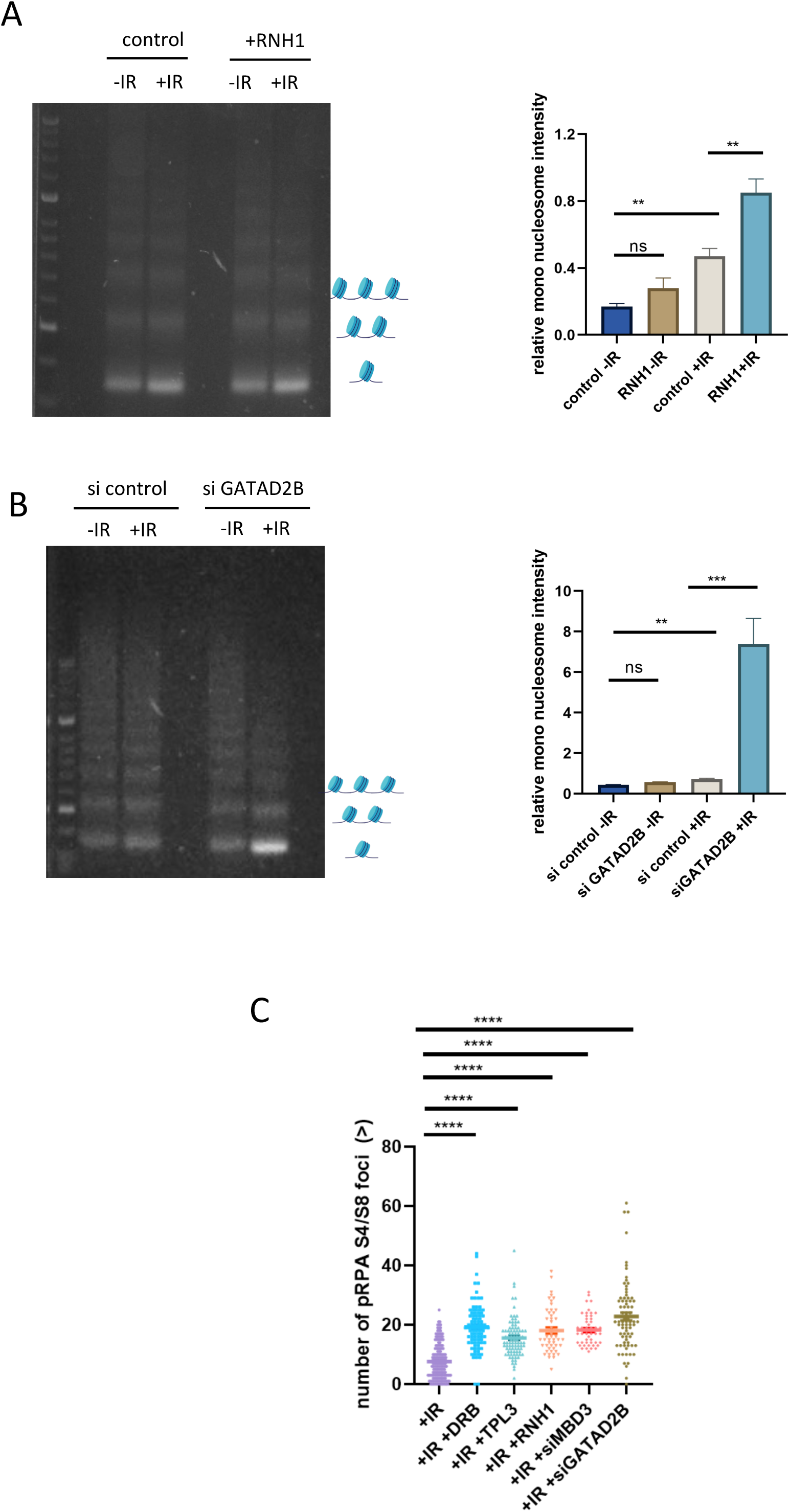

**Expanded View Figure 12.**
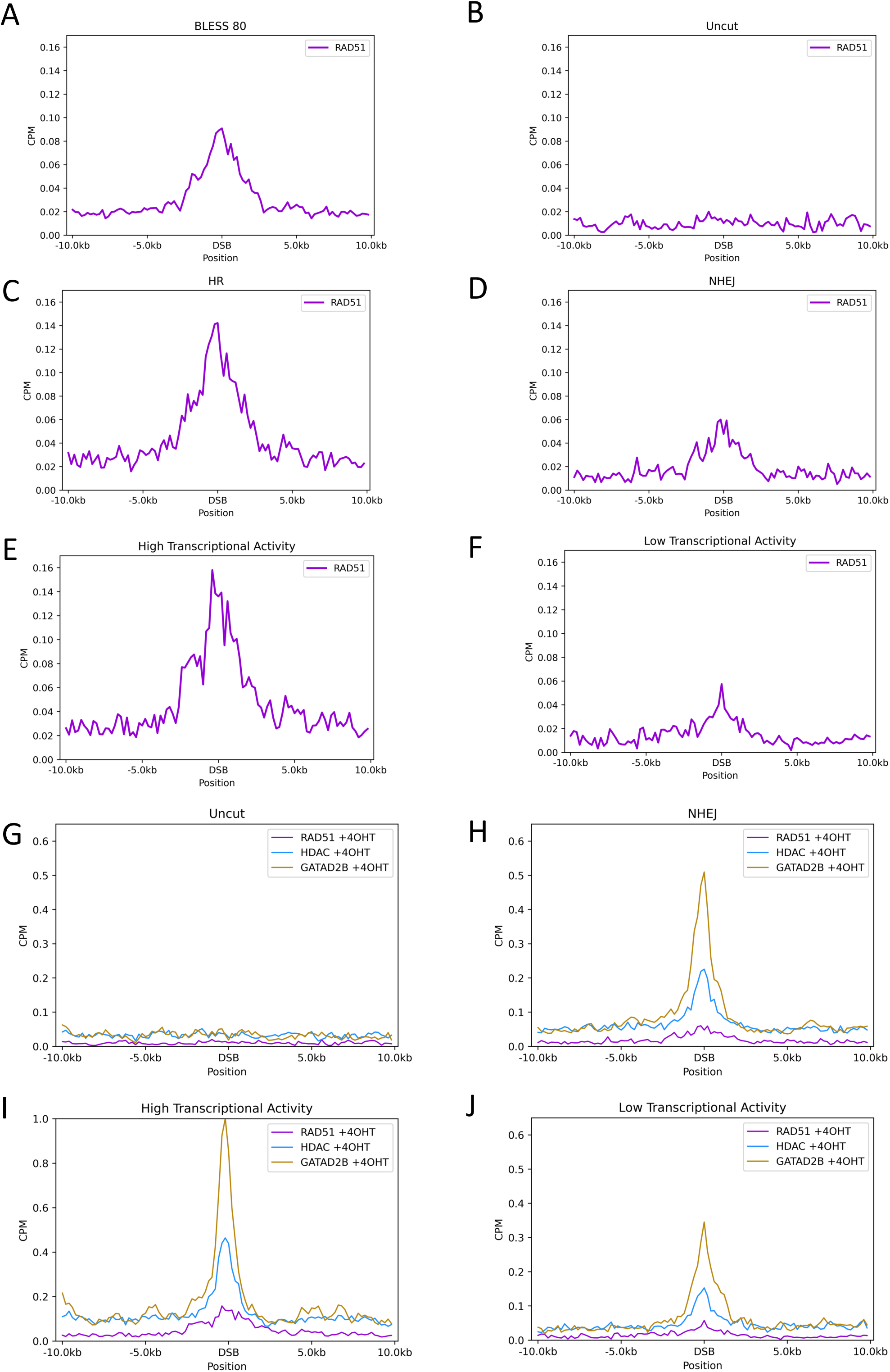

**Expanded View Figure 13.**
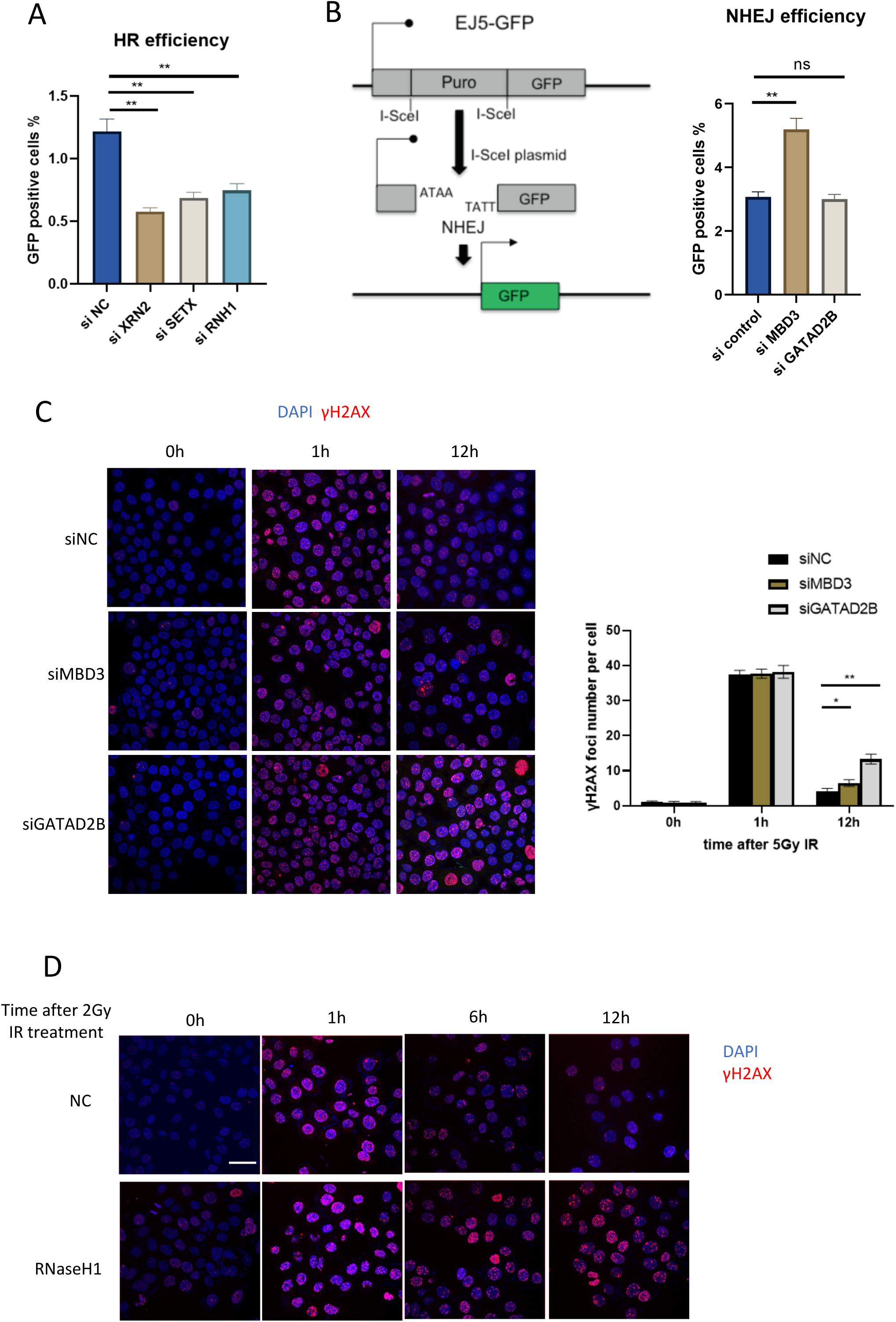

