## Supplementary Figure legends for "GATAD2B containing NuRD complex drives R-loop dependent chromatin boundary formation at double strand breaks"

### Expanded View Figure Legends

#### Fig EV1

A) Silver-stained gel showing proteins in S9.6 IP and no antibody (B) control.

B) Western blot analysis of γH2AX showing that DSBs were successfully induced by 10 Gy IR.

C) Western blot analysis showing presence of XRN2, DHX9 and H3, in S9.6 IP and control (B) samples.

D) Silver-stained gel showing proteins in S9.6 IP samples prepared for mass spectroscopy analysis.

E) Western blot analysis showing presence of XRN2, DHX9 and H3 in S9.6 IP samples prepared for mass spectroscopy analysis.

F) Table showing known R-loop binding factors which have been verified by mass spectroscopy in S9.6 IP in no damage condition and their intensities.

G) Table showing proteins which have been identified by mass spectroscopy to preferentially interact with R-loops upon IR treatment.

#### Fig EV2

A) Representative confocal images showing single antibody controls for PLA showed in Figure 1.

B) Left: Representative confocal images showing PLA of GATAD2B and S9.6 in cells with or without IR and treatment with PARP1 or ATM/ATR inhibitors. Right: quantification of left, error bar = mean ± SEM, significance was determined using non-parametric Mann-Whitney test. ****$p \leq 0.0001$.

C) Representative confocal images showing PLA of HDAC1 and S9.6 in cells with or without IR and overexpression of RNAseH1 Left: representative confocal microscopy images and single antibody PLA control; right: quantification of left, error bar = mean ± SEM, significance was determined using non-parametric Mann-Whitney test. ****$p \leq 0.0001$.

#### Fig EV3

A) Laser stripping of MBD3-GFP cells with or without treatment with transcription inhibitors triptolide (TPL3) and DRB. Representative spinning disk confocal microscopy images and quantification (n≥10) showing GFP signals before and after laser striping at indicated time points; error bar = mean ± SEM.

B) Laser stripping of MBD3-GFP cells with or without transiently expression of RNAseH1-RFP plasmid. Representative spinning disk confocal microscopy images and quantification (n ≥10) showing GFP and RFP signals before and after laser striping at indicated time points; error bar = mean ± SEM.

C) Quantification (n≥10) showing GATAD2B-GFP and RNH1-mCherry signals before and after laser striping at indicated time points; error bar = mean ± SEM.

D) ChIP-qPCR bar charts showing levels of HDAC1 (left) and GATAD2B (right) at three genes known to be bound by NuRD complex in non-damage condition in cells treated with Triptolide or overexpressing RNaseH1.

**Fig EV4**

A) Representative confocal images showing PLA of MBD3 and γH2AX in cells with or without IR and transcription inhibition (TLP3 or DRB) or overexpression of RNAseH1. IR=10Gy. Left: representative confocal microscopy images; right: quantification of left, error bar = mean ± SEM, significance was determined using non-parametric Mann-Whitney test. ****$p \leq 0.0001$.

B) Western blot showing efficiency of siRNA mediated knockdown of GATAD2B and MBD3, NC = negative control. Tubulin was used as loading control.

C) Representative confocal images showing PLA of HDAC1 and γH2AX in cells with or without IR and depleted of ZMYND8 or KDM5A, IR=10Gy. Left: representative confocal microscopy images; right: quantification of left, error bar = mean ± SEM, significance was determined using non-parametric Mann-Whitney test. ****$p \leq 0.0001$.

**Fig EV5**

A) PCA plots showing input (control), GATAD2B and HDAC1 ChIP-seq replicates comparing consensus peaks detected using MACS2.

B) Metagene profile showing log2fold (+4OHT/-4OHT) ChIP-seq enrichment of GATAD2B, HDAC1 and input at AsiSI cut sites (as defined by BLESS technique).

**Fig EV6**

A) Metagene profile showing ChIP-seq enrichment of GATAD2B, HDAC1 and R-loops at AsiSI uncut sites (as defined by BLESS technique).

B) Heatmaps representing GATAD2B and HDAC1 ChIP-seq count over a 20 kb window centered on the DSB before(−4OHT) and after(+4OHT) DSB induction. DSBs are sorted according to decreasing cutting efficiency.

**Fig EV7**

A) Metagene profile showing ChIP-seq enrichment of GATAD2B, HDAC1 and R-loops at AsiSI cut sites in low transcribed regions.

B) Box plot graph showing levels of HDAC1 and GATAD2B ChIP-seq reads mapping to AsiSI sites in presence or absence of 4OHT at high transcription regions.

C) Box plot graph showing levels of HDAC1 and GATAD2B ChIP-seq reads mapping to AsiSI sites in presence or absence of 4OHT at low transcription regions.

D) Metagene profile showing ChIP-seq enrichment of GATAD2B, HDAC1 and R-loops at NHEJ prone AsiSI cut sites (as defined by BLESS technique).

E) Box plot graph showing levels of HDAC1 and GATAD2B ChIP-seq reads mapping to AsiSI sites in presence or absence of 4OHT at HR prone regions.

F) Box plot graph showing levels of HDAC1 and GATAD2B ChIP-seq reads mapping to AsiSI sites in presence or absence of 4OHT at NHEJ prone regions.

**Fig EV8**

A) Heatmaps representing GATAD2B and HDAC1 ChIP-seq, DRIP-seq and H4K12Ac ChIP-seq count over a 20 kb window centered on the DSB before(−4OHT) and after(+4OHT) DSB induction. DSBs are sorted according to decreasing cutting efficiency.

B) Metagene profile showing ChIP-seq enrichment of GATAD2B, HDAC1 and H4K12ac at AsiSI cut sites in low transcribed regions.

C) Metagene profile showing ChIP-seq enrichment of GATAD2B, HDAC1 and H4K12ac at NHEJ prone AsiSI cut sites.

**Fig EV9**

A-C) IGV screenshots showing HDAC1 and GATAD2B ChIP-Seq, DRIP-seq and H4K12Ac ChIP-seq reads count in no damage (-4OHT) and damage (+4OHT) conditions at representative AsiSI cutting sites.

**Fig EV10**

A) Left: representative confocal images showing PLA of pan-acetyl H3 and γH2AX in cells with or without IR, and overexpression of RNAseH1 or depletion of GATAD2B and MBD3. IR=10Gy. Right: quantification of left, error bar = mean ± SEM, significance was determined using non-parametric Mann-Whitney test. ****$p \leq 0.0001$

B) Bar chart showing relative pan-acetyl H4 and H4 ChIP levels at the DS1 loci next to AsiSI cut in cells with or without 4OHT and depletion of GATAD2B, error bar = mean ± SEM, significance was determined using non-parametric Mann-Whitney test. ***$p \leq 0.001$, **$p \leq 0.01$,*$p \leq 0.05$, n.s. not significant.

C) Bar chart showing relative pan-acetyl H4 and H4 ChIP levels at the DS2 loci next to AsiSI cut in cells with or without 4OHT and depletion of GATAD2B, error bar = mean ± SEM, significance was determined using non-parametric Mann-Whitney test. ***$p \leq 0.001$, **$p \leq 0.01$,*$p \leq 0.05$, n.s. not significant.

D) Western blot showing the levels of Pan H4-Ac, GATAD2B and H4 in control (siNC) and siGATAD2B cells.

**Fig EV11**

A) Left: DNA gel showing nucleosome profile of wt and cells overexpressing RNaseH after MNase treatment and DNA extraction. Right: Bar chart showing relative mono-nucleosome intensity of each lane. **$p \leq 0.01$, n.s. not significant.

B) Left: DNA gel showing nucleosome profile of wt and cells depleted of GATAD2B after MNase treatment and DNA extraction. Right:Bar chart showing relative mono nucleosome intensity of each lane. ***$p \leq 0.001$, **$p \leq 0.01$, n.s. not significant.

C) Graph showing number of pRPA foci that were quntified at prominence set to 2500 in FIJI software.

**Fig EV12**

A) Metagene profile showing ChIP-seq enrichment of RAD51 at BLESS 80 AsiSI cut sites.

B) Metagene profile showing ChIP-seq enrichment of RAD51 at uncut AsiSI sites.

C) Metagene profile showing ChIP-seq enrichment of RAD51 at cut HR prone AsiSI sites.
D) Metagene profile showing ChIP-seq enrichment of RAD51 at cut NHEJ prone AsiSI sites.
E) Metagene profile showing ChIP-seq enrichment of RAD51 at highly transcribed cut AsiSI sites.
F) Metagene profile showing ChIP-seq enrichment of RAD51 at low transcribed cut AsiSI sites.
G) Metagene profile showing ChIP-seq enrichment of GATAD2B, HDAC1 and RAD51 at uncut AsiSI sites.
H) Metagene profile showing ChIP-seq enrichment of GATAD2B, HDAC1 and RAD51 at NHEJ prone cut AsiSI sites.
I) Metagene profile showing ChIP-seq enrichment of GATAD2B, HDAC1 and RAD51 at highly transcribed cut AsiSI sites.
J) Metagene profile showing ChIP-seq enrichment of GATAD2B, HDAC1 and RAD51 at low transcribed cut AsiSI sites.

**Fig EV13**

A) Bar charts showing HR repair efficiency under DSB inducted by I-SceI in U2OS cells containing HR reporter, and control (siNC) or knock down of XRN2, SETX or with overexpression of RNH1. error bar = mean ± SEM, significance was determined using non-parametric Mann-Whitney test. ** $p \le 0.01$.

B) Left: drawing indicating structure of NHEJ reporter cassette. Right: Bar charts showing NHEJ repair efficiency under DSB inducted by I-SceI in U2OS cells containing NHEJ reporter, and knock down of MBD3 and GATAD2B

C) Immunofluorescence of γH2AX in cells with IR, and depletion of GATAD2B and MBD3. Left: representative confocal microscopy images; right: quantification of left, error bar = mean ± SEM, significance was determined using non-parametric Mann-Whitney test. ***$p \le 0.001$, ** $p \le 0.01$, * $p \le 0.05$.

D) Immunofluorescence of γH2AX in cells with IR, and overexpression of RNaseH.

Figure S9. The Model of chromatin boundary around DSBs established by GATAD2B-NuRD complex and R-loops.
