## Supplementary Tables for "GATAD2B containing NuRD complex drives R-loop dependent chromatin boundary formation at double strand breaks"

**Supplementary material**

**Supplementary Table**

| **Reagent or Resource** | **Source** | **Identifier** |
|---|---|---|
| **Antibodies** | | |
| Rabbit-Anti-GATAD2B | Abcam | Ab76925 |
| Rabbit-Anti-MBD3 | Abcam | Ab157464 |
| Mouse Anti-HDAC1 | Santa Cruz | Sc-81598 |
| Anti-phospho-Histone H2A.X (Ser139), clone JBW301 | Merck Millipore | 05-636 |
| Anti-Phospho-Histone H2A.X (Ser139) (20E3) Rabbit mAb | Cell Signaling Technology | 9718S |
| Anti-DNA-RNA Hybrid Antibody, clone S9.6, mouse monoclonal | Merck | MABE1095 |
| Anti-DNA–RNA hybrid antibody, clone S9.6, rabbit monoclonal | Absolute Antibody | Ab01137-23.0 |
| Histone H3ac (pan-acetyl) antibody, Rabbit polyclonal | Active Motif | 39140 |
| Histone H4ac (pan-acetyl) antibody, Rabbit polyclonal | Active Motif | 39043 |
| Anti-Histone H3 | Abcam | Ab1791 |
| Anti-Histone H4 Polyclonal antibody | Proteintech | 16047-1-AP |
| Anti-beta Tubulin antibody | Abcam | Ab6046 |
| Anti-RNA polymerase II CTD repeat YSPTSPS antibody | Abcam | Ab26721 |
| Anti-RNA polymerase II CTD repeat YSPTSPS (phospho S2) antibody | Abcam | Ab5095 |
| Anti- XRN2 Antibody (H-3) | Santa Cruz | sc-365258 |
| Anti-BrdU antibody, Mouse monoclonal | Sigma | B-8434 |
| Anti-Rad51 Antibody | Abcam | Ab63801 |

| | | |
|---|---|---|
| Anti-V5 tag antibody [SV5-P-K] | Abcam | Ab206566 |
| Donkey Anti-Rabbit IgG (H+L) Highly Cross-Adsorbed Secondary Antibody, Alexa Fluor™ 488 | Thermo Fisher | A-21206 |
| Donkey anti-Mouse IgG (H+L) Highly Cross-Adsorbed Secondary Antibody, Alexa Fluor™ 555 | Thermo Fisher | A-31570 |
| Rabbit anti-Phospho RPA32 (S4/S8) Antibody | Bethyl Laboratories | A300-245A |
| Anti-RNA Helicase A (DHX9) | Abcam | ab26271 |
| **Chemicals, peptides, and recombinant proteins** | | |
| Triptolide | Cayman Chemical | CAY11973 |
| DRB | Cayman | 10010302 |
| Olaparib | Cayman | CAY-10621 |
| 4-OHT | Cayman | CAY-14854 |
| 5-Bromo-2’-deoxyuridine (5-BrDU) | Sigma | B-5002 |
| RNaseH | New England Biolabs | M0297S |
| Lipofectamine 3000 | Thermo Fisher | L3000008 |
| Lipofectamine RNAiMAX | Thermo Fisher | 13778075 |
| Hoechst 33342 | Thermo Fisher | R37605 |
| Dynabeads Protein A | Thermo Fisher | 10002D |
| Micrococcal Nuclease | New England Biolabs | M0247S |
| Dynabeads Protein G | Thermo Fisher | 10004D |
| AZD5305 | Cayman | 2589531-76-8 |
| Ku-55933 | Cayman | 587871-26-9 |
| AZ20 | Selleckchem | S7050 |
| **Critical commercial assays** | | |
| Duolink In Situ Red Starter Kit Mouse/Rabbit | Sigma | DUO92101-1KT |
| CometAssay Electrophoresis Starter | Bio-Techne | 4250-050-ESK |

| | | |
|---|---|---|
| Kit | | |
| Experimental models: Cell lines | | |
| HeLa | ATCC | CCL-2 |
| U-2 OS cells | ATCC | HTB-96 |
| HEK293T cells | ATCC | CRL-3216 |
| U-2 OS DR-GFP cells | A gift from Xingzhi Xu | N/A |
| U-2 OS EJ5-GFP cells | A gift from Xingzhi Xu | N/A |
| AsiSI-ER U2-OS | A gift from Gaelle Legube | N/A |
| Oligonucleotides | | |
| SiGATAD2B siGENOME SMARTPOOL | Dharmacon | M-013892-01-0010 |
| SiMBD3 siGENOME SMARTPOOL | Dharmacon | M-013616-01-0010 |
| ON-TARGETplus Non-targeting Control | Dharmacon | D-001810-01-20 |
| SiXRN2 siGENOME SMARTPOOL | Dharmacon | M-017622-01-0005 |
| SiSETX siGENOME SMARTPOOL | Dharmacon | M-021420-00-0005 |
| SiZMYND8 siGENOME SMARTPOOL | Dharmacon | M-017354-00-0005 |
| SiKDM5A siGENOME SMARTPOOL | Dharmacon | M-003297-02-0005 |
| SiRNase H1 siGENOME SMARTPOOL | Dharmacon | M-012595-01-0005 |

| | | |
|---|---|---|
| **Recombinant DNA** | | |
| ppyCAG_RNaseH1_WT | Addgene | #111905 |
| pCMV-mCherry-RNaseH1 | This Study | N/A |
| pCMV-GFP-MBD3 | This Study | N/A |
| pCMV-GFP-GATAD2B | This Study | N/A |
| pH2B-PAGFP | (8) | N/A |
| peGFP-BZIP | | N/A |

| Software and algorithms | | |
|---|---|---|
| GraphPad Prism 9 | www.graphpad.com | N/A |
| Fiji | (9) | N/A |
| CellProfiler | (10) | N/A |
| Flowjo | www.Flowjo.com | N/A |
| bedtools multicov | https://bedtools.readthedocs.io/ | N/A |

Primers applied in ChIP-qPCR and RT-qPCR

| **Name** | **Sequences (5'-3')** |
|---|---|
| **28s-F** | **TTCCCTCCGAAGTTTCCCTC** |
| **28s-R** | **ACTAGGCACTCGCATTCCAC** |
| **DS2 +0.3-F** | **CCAGCAGTAAAGGGGAGACAGA** |
| **DS2 +0.3-R** | **CTGTTCAATCGTCTGCCCTTC** |
| **DS2 +1.8-F** | **GAAGCCATCCTACTCTTCTCACCT** |
| **DS2 +1.8-R** | **GCTGGAGATGATGAAGCCCA** |
| **DS2 +3.6-F** | **GCCCAGCTAAGATCTTCCTTCA** |
| **DS2 +3.6-R** | **CTCCTTTGCCCTGAGAAGTGA** |
| **DS1 +0.3-F** | **AGGACTGGTTTGCAAGGATG** |
| **DS1 +0.3-R** | **ACCCCCATCTCAAATGACAA** |
| **DS1 +1.0-F** | **AGGAATTGACTGCGGTGTTC** |
| **DS1 +1.0-R** | **GGGGAGGAGGAAAGGTGTAG** |
| **DS1 +2.0-F** | **GCCATAACAGAGGGTGGAAA** |
| **DS1 +2.0-R** | **AACTTTAGGATGGGGCTGCT** |
| **DS1+3.0-F** | **TGTAGCCACAGTTTGCCTGT** |
| **DS1+3.0-R** | **CTCCTCTATTGTCACCTGGAAGAC** |
| **DS1+5.0-F** | **GGCCAACATCCCTGATGACTAC** |
| **DS1+5.0-R** | **CACCCTTGCCAGCATTTGTT** |
| **MACROD2 pro-F** | **ACGCAGCACAGTCCTTTGG** |
| **MACROD2 pro-R** | **AGGACCTGAATTCTGTGGTGG** |
| **GAPDH PRO-F** | **ATCCAAGCGTGTAAGGGTCC** |
| **GAPDH PRO-R** | **TAGGGGGGAAGGGACTGAGA** |
